# Structural basis of the T4 bacteriophage primosome assembly and primer synthesis

**DOI:** 10.1101/2023.05.03.539249

**Authors:** Xiang Feng, Michelle M. Spiering, Ruda de Luna Almeida Santos, Stephen J. Benkovic, Huilin Li

## Abstract

The T4 bacteriophage gp41 helicase and gp61 primase assemble into a primosome complex to couple DNA unwinding with RNA primer synthesis for DNA replication. How a primosome is assembled and how the length of the RNA primer is defined in the T4 bacteriophage, or in any model system, are unclear. Here we report a series of cryo-EM structures of T4 primosome assembly intermediates at resolutions up to 2.7 Å. We show that the gp41 helicase is an open spiral in the absence of ssDNA, and ssDNA binding triggers a large-scale scissor-like conformational change that drives the open spiral to a closed ring that activates the helicase. We found that the activation of the gp41 helicase exposes a cryptic hydrophobic primase-binding surface allowing for the recruitment of the gp61 primase. The primase binds the gp41 helicase in a bipartite mode in which the N-terminal Zn-binding domain (ZBD) and the C-terminal RNA polymerase domain (RPD) each contain a helicase-interacting motif (HIM1 and HIM2, respectively) that bind to separate gp41 N-terminal hairpin dimers, leading to the assembly of one primase on the helicase hexamer. Based on two observed primosome conformations – one in a DNA-scanning mode and the other in a post RNA primer-synthesis mode – we suggest that the linker loop between the gp61 ZBD and RPD contributes to the T4 pentaribonucleotide primer. Our study reveals T4 primosome assembly process and sheds light on RNA primer synthesis mechanism.

## INTRODUCTION

During DNA replication, the double-stranded DNA (dsDNA) is unwound by a ring-shaped helicase, which moves in the 3’ to 5’ direction on the leading-strand DNA in eukaryotes and in the 5’ to 3’ direction on the lagging-strand DNA in prokaryotes and bacteriophages^1,2^. Because all DNA polymerases extend only the 3’-OH of an existing oligonucleotide, an RNA primer is synthesized to initiate DNA synthesis. Primers are short RNA oligonucleotides synthesized by primases and are complementary to the parent DNA template. Primases are RNA polymerases in bacteria and bacteriophage, but are dual-functioning RNA and DNA polymerases in eukaryotes (e.g. - Pol α)^3,4^. The priming reaction occurs frequently on the lagging-strand DNA in order to start each Okazaki fragment. The eukaryotic Pol α is associated with the replicative CMG helicase indirectly via the Ctf4 trimer^1,5^ and synthesizes an RNA-DNA hybrid of 20 nucleotides (nt). In bacteria and phages, the primase is directly associated with the helicase and synthesizes RNA primers of various length, e.g.- ∼11 nt by the bacterial DnaG primase, 4 nt by the T7 gp4 primase, and 5 nt by the T4 gp61 primase. The priming mechanism is perhaps the least well understood aspect in DNA replication in all domains of life^3,4,6-8^. The primase counting or measuring capability to determine the length of the primer synthesized has been discussed hypothetically for the archaeal and eukaryotic primase complexes^9-11^.

In bacteria and phages, the helicase and primase assemble into a functional complex known as a primosome to couple DNA unwinding with RNA primer synthesis. The bacterial DnaB helicase is composed of an N-terminal domain (NTD) and a C-terminal domain (CTD). The DnaB helicase assembles into a two-tiered hexameric ring where the N-tier has three-fold symmetry due to dimerization of the N-terminal helical hairpins, while the C-tier has nearly six-fold symmetry composed of six CTDs with six ATP binding pockets at the CTD-CTD interfaces^12^; and the N-tier and the C-tier are staggered in a domain-swapped fashion^12,13^. The DnaB helicase was observed in two states: a “dilated” state where three N-terminal helical hairpin dimers form an equal-sided triangle and the C-tier encircles a large channel, and a “constricted” lock-washer state where the N-terminal helical hairpin dimers swivel inward to bring the CTDs towards each other constricting the central chamber to accommodate only single-stranded DNA (ssDNA)^14,15^. The bacterial DnaB helicase is thought to translocate on ssDNA utilizing a “rotary-staircase” mechanism in which subunits translocate sequentially from one end of the “lock washer” to the next at the expense of ATP hydrolysis^1,13,15^. The bacterial DnaG primase is a 65-kDa, three-domain protein with an N-terminal Zn-binding domain (ZBD), a core RNA polymerase domain (RPD), and a C-terminal helicase-interacting domain (HID)^16^. Structures are available for the individual domains, but not for any full-length primase, suggesting that these domains are flexibly connected^17-22^. It’s been suggested that the DnaG primase and DnaB helicase form a primosome with a 3:6 stoichiometry mediated by the interaction between the C-terminal HID of DnaG and the N-terminal hairpin dimer of DnaB^17,23^. However, no structure has been reported for any bacterial primosome – either alone or in complex with ssDNA and/or an RNA primer.

The T7 bacteriophage gp4 helicase belongs to the superfamily of prokaryotic replicative helicases that includes DnaB^2^. The T7 replication system is unique in that the primase and the helicase are fused into a single polypeptide, gp4, such that the stoichiometry of the T7 primosome must be 6 primases to 6 helicases. The gp4 helicase region lacks the N-tier ring and has only the C-tier ring equivalent to the bacterial DnaB helicase. Therefore, the T7 primosome has largely a six-fold symmetrical architecture. A recent cryo-EM study revealed that the T7 gp4 helicase has a right-handed spiral shape and exists in several states, suggesting that the helicase translocates on the lagging-strand DNA with a hand-over-hand “rotary-staircase” mechanism^24^. The same study also captured a state in which the ZBD of the gp4 primase region was handing over the RNA primer to the DNA polymerase. This structure revealed the coordination between the primase and the DNA polymerase but does not inform on how the primer was synthesized.

T4 bacteriophage has been a classic model system for studying DNA replication and RNA priming mechanisms^6,25^. The T4 gp41 helicase also belongs to the prokaryotic replicative helicase superfamily^2^. The gp41 helicase and the gp61 primase assemble into a primosome complex to couple parental DNA unwinding with priming activity on the lagging-strand DNA^25^. The domain architecture of the gp41 helicase resembles the bacterial DnaB helicase^13,14^ with an NTD and a RecA-like CTD that are connected by a linking helix (LH) (**Fig. 1a**). In the presence of ATP, the gp41 helicase monomers assemble into a hexamer to encircle the lagging-strand DNA^26^ and move in a 5’ to 3’ direction to unwind duplex DNA at a rate of 30 bp/s^27,28^. The gp61 primase is composed of an N-terminal ZBD and a C-terminal RPD that are linked by an 18-residue flexible loop (**Fig. 1a**). The gp61 primase recognizes either 5′-GTT or 5′-GCT on the ssDNA template as a priming start site and synthesizes a pentaribonucleotide primer^29^. The underlying mechanism for synthesizing this length of primer by the T4 primosome is unknown.

**Figure 1.**
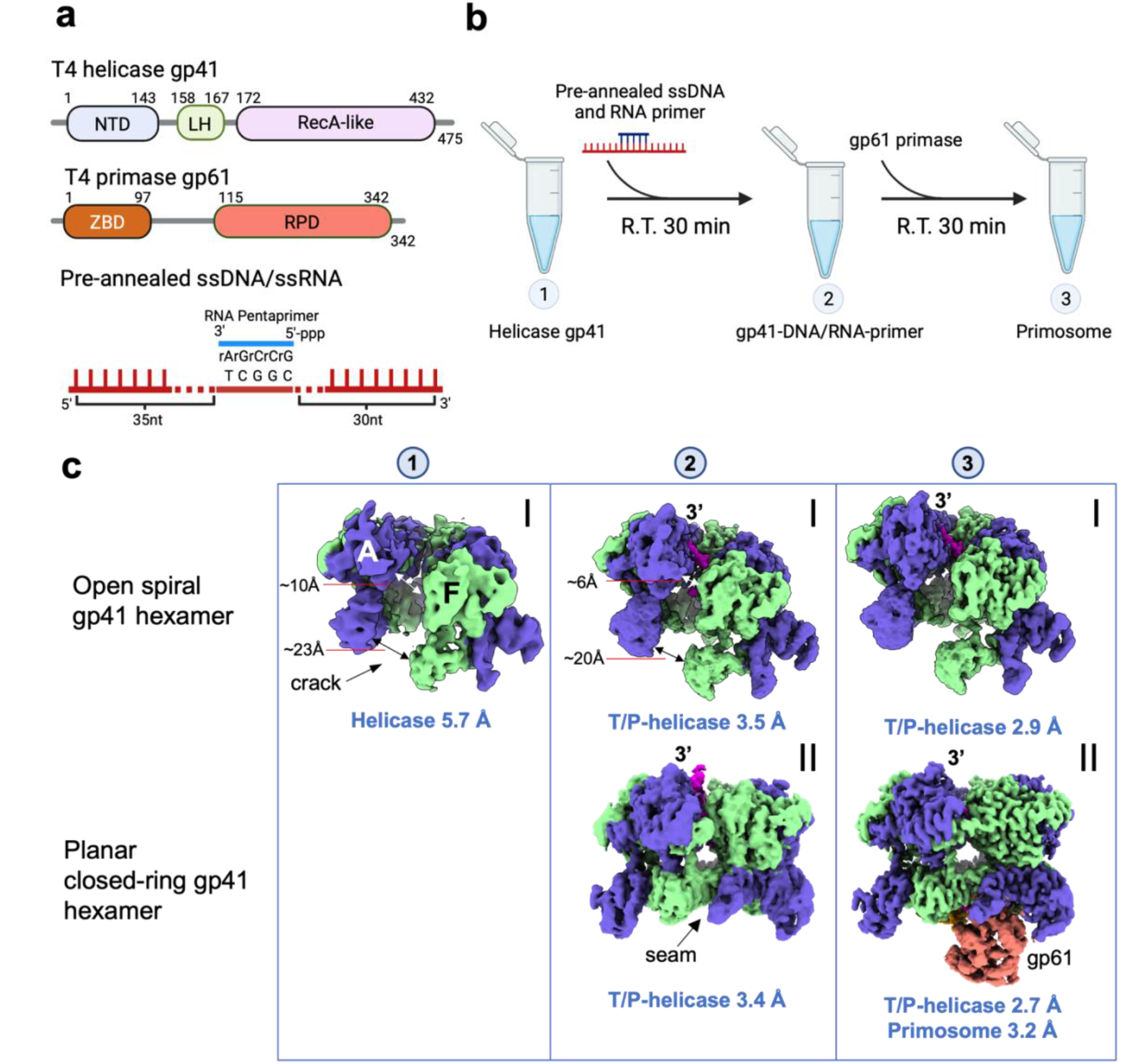
In vitro assembly and cryo-EM analysis of the T4 primosome. **a)**Domain architecture of the gp41 helicase, the gp61 primase, and the ssDNA/RNA primer construct used in the in vitro assembling of the primosome. **b**) Step-by-step process used to assemble the T4 primosome. **c**) 3D EM maps of the step 1 complex – (I) the hexameric gp41 helicase in an open spiral configuration; two step 2 complexes – the ssDNA-bound gp41 helicase in the (I) open spiral and (II) planar closed-ring configurations; and two step 3 complexes – (I) the ssDNA-bound gp41 helicase in an open spiral form and (II) the primosome. The average resolution of each map is labeled below the respective map. The primosome map (3-II) is a composite map of the separately refined maps of the ssDNA-bound gp41 helicase (2.7 Å) and the gp61 primase region of ssDNA-bound gp41 helicase-gp61 primase complex. The overall resolution of the composite primosome map reaches 3.2 Å, but the resolution of primase region is lower; see the detailed local resolution estimation in **Supplementary Fig. 5**. The EM maps are postprocessed by DeepEMhancer (using the “tightTarget” model) and aligned in such way that the gap in open spirals or the seam in closed rings face the readers. The two open-spiral, ssDNA-bound gp41 helicase structures (2-I and 3-I) are the same; and the closed-ring gp41 helicase structure (2-II) is essentially the same as the gp41 helicase region in the primosome structure (3-II).

In this study, we determined a series of structures of T4 primosome assembly intermediates and the assembled primosome in a DNA-scanning mode and in a post RNA primer-synthesis mode. We show that one gp61 primase binds to the helicase hexamer through interacting interfaces on both the ZBD and RPD of the primase. We also explored the influence the primase linker loop has over the length of the synthesized RNA primer. In sum, the structural study has enabled us to propose a unique helicase activation mechanism, a detailed process for primosome assembly, and a plausible mechanism for pentaribonucleotide primer synthesis by the T4 primosome.

## RESULTS

### Cryo-EM of T4 primosome assembly intermediates

We assembled the T4 primosome in vitro with purified components through a process like the one described previously^30^ (**Fig. 1b**) and resolved the intermediate states during primosome assembly (**Fig. 1c**). We started with the assembly of the gp41 helicase hexamer by adding ATPγS (step 1). Our previous work showed that a 45-nt ssDNA with a priming recognition site was required for detectable priming activity ^31^ and that a longer ssDNA substrate might be required to accommodate a DNA loop between the helicase and primase ^32^. For these reasons, we designed a 70-nt ssDNA template with a 5-nt RNA primer annealed in the middle as a substrate for the helicase and product for the primase and added to the helicase solution (step 2). Finally, we added the purified gp61 primase to form a minimalist T4 primosome (step 3). Each mixture was incubated for 30 min at room temperature before the reaction products were withdrawn to prepare the cryo-EM grids. Through cryo-EM analysis, we determined a total of eight 3D maps that represented three states of the primosome assembly and/or functional states (**Fig. 1c, Supplementary Fig. 1-6**).

The 3D map (map 1-I at 5.7 Å resolution) derived from step 1 showed the gp41 helicase hexamer in the absence of ssDNA as a right-handed open spiral with a 10-Å gap between subunits A and F wide enough for the passage of an ssDNA. From step 2, we derived two 3D maps of the gp41 helicase hexamer bound to the ssDNA template. One map (map 2-I at 3.5 Å resolution) showed the ssDNA template in the helicase central channel, but the helicase hexamer remained in the right-handed open spiral form. In contrast, the other map (map 2-II at 3.4 Å resolution) showed the gp41 helicase bound to the ssDNA template that had undergone major conformational changes to become a planar ring closing the gap between subunits A and F and likely represents the active helicase configuration. From step 3, we determined two additional 3D maps. One map (map 3-I at 2.9 Å resolution) showed an open spiral gp41 helicase hexamer with the ssDNA template bound inside the central channel similar to map 2-I. The other map (map 3-II at 3.2 Å overall resolution) showed the active gp41 helicase hexamer as a planar ring encircling the ssDNA template (2.7 Å local resolution) with a gp61 primase bound. The high resolution achieved for each map allowed for atomic modeling of the intermediate states during primosome assembly.

### DNA template binding induces conformational changes that activate the helicase

The first state of primosome assembly (map 1-I at 5.7 Å resolution) showed the gp41 helicase in the absence of ssDNA as a two-tiered asymmetric homo-hexamer with an N-tier and a C-tier (**Fig. 1c**). The NTD can be further divided into a globular subdomain followed by a helical hairpin. The NTDs of two neighboring subunits form an antiparallel dimer via angled packing of the two helical hairpins. Three such head-to-tail dimers associate mediated by interactions between the N-terminal globular subdomains leading to the triangular appearance of the N-tier and the trimer-of-dimers architecture of the gp41 helicase. In the C-tier of the hexameric helicase, the six CTDs are arranged nearly symmetrically with five nucleotide binding pockets at the subunit interfaces occupied with ATPγS molecules. The gp41 helicase forms a right-handed open spiral with a 10 Å gap between subunits A and F. In the absence of ssDNA, subunits A and F that line the gap of the open spiral were partially mobile to perhaps facilitate the entry or exit of ssDNA from the central channel.

The second state of primosome assembly is represented by map 2-I at 3.5 Å resolution and map 3-I at 2.9 Å, each with the ssDNA template bound in the central channel but with the gp41 helicase remaining an open spiral. Initial binding of the ssDNA caused only minor changes to the open spiral structure making it slightly more compact with a narrower gap and less mobility in subunits A and F (**Supplementary Fig. 1e**). These minimal conformational changes indicate that the DNA-free gp41 helicase is competent for loading onto ssDNA, although a gp59 helicase loader protein is necessary for efficient helicase loading in vivo. This is very different from the DnaB helicase, which has a closed, planar conformation in the nucleotide-bound, ssDNA-free state necessitating a helicase loader (the DnaC hexamer^12,15^ or loaders from bacteriophage^33^) to open the DnaB helicase ring to load onto ssDNA.

In contrast, the third state of primosome assembly is represented by map 2-II at 3.4 Å resolution and map 3-II at 2.7 Å resolution. Both showed that three major conformational changes occurred when the ssDNA template-bound, open spiral of the gp41 helicase transitioned to a planar, closed-ring state (**Fig. 2a-b**). First, the N-tier rotated in plane by 60° with respect to the C-tier leading to a domain-swapped packing in the closed state, i.e. - each CTD now aligned with the neighboring NTD. The domain-swapped arrangement has also been observed in other members of the prokaryotic replicative helicase superfamily^12,13,15,23,34-36^. Second, the linking helix (LH) from the N-tier relocated to interact with the CTD of the neighboring subunit. Finally, the N-terminal helical hairpins (enclosed in a yellow, dashed-line shape) closed in a scissor-like motion that increased their crossing angle from 100° in the open spiral form to 165° and almost antiparallel in the closed-ring form (**Fig. 2a-d**). Because the gap between subunits A and F is now closed, this map likely represents the active gp41 helicase configuration. This scissor-like activation mechanism appears to be unique to the gp41 helicase. The *E. coli* DnaB helicase was reported to use a very different aperture-like motion to close the hexameric ring and activate the DnaB helicase^15^ (**Fig.2 e-f**).

**Figure 2.**
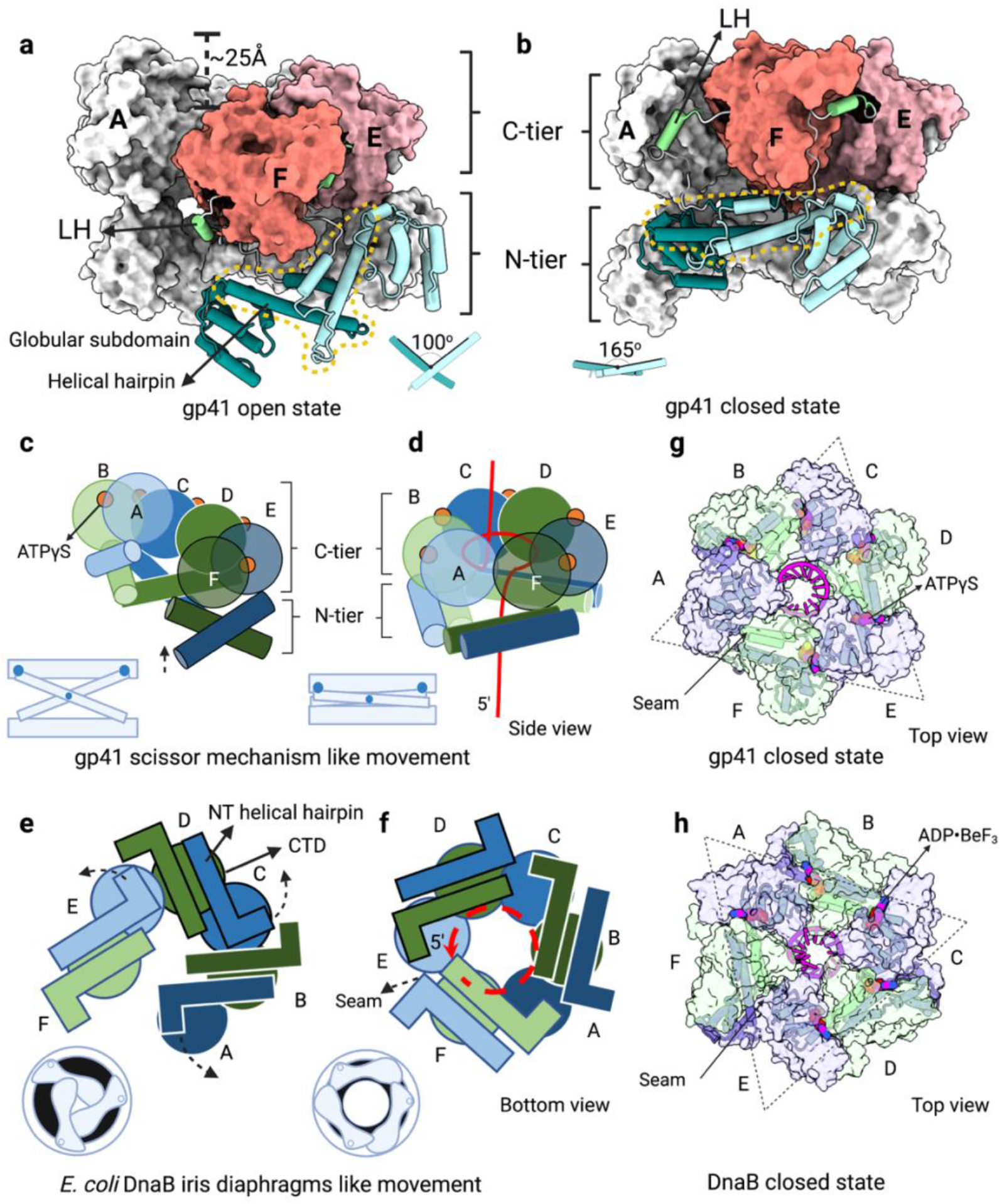
Conformational changes from the inactive, open-spiral to the active, closed-ring structures of the ssDNA-bound gp41 helicase. Side views of **a)** the open spiral and **b)** the closed-ring structure. The domains are colored as in Fig.1a. Subunits A-D are shown as grey surfaces. The NTDs and LHs of subunits E and F are in cartoon form and their CTD are shown in colored surfaces. Illustrations depict the scissor-like motion of the gp41 helical hairpins in **c)** the inactive, open spiral and **d)** the active, closed-ring structures. In contrast, illustrations depict the aperture-like swivel motion of the N-terminal helical hairpins of the *E. coli* DnaB helicase in **e)** the inactive, open spiral and **f)** the active, closed ring (PDB entries 6QEL and 6QEM). The red curves in d) and f) represent the lagging-strand DNA. Top surface views of **g)** the gp41 helicase showing the bound ssDNA and ATPγS molecules (this study) and **h)** the DnaB helicase showing the bound ssDNA and nucleotides (PDB entry 6QEM). The dashed triangles indicate the interdimeric and intradimeric location of the seam in **g)** the T4 helicase and **h)** the DnaB helicase, respectively.

### Active T4 helicase is a planar ring with DNA-translocating loops spiraling around the central channel

In the active gp41 helicase, five ATPγS molecules were resolved in the C-tier with the nucleotide-binding pocket between the CTDs of subunits A and F unoccupied (**Fig. 2g**). This is comparable to the five ADP-BeF_3_ molecules observed in the DNA-bound DnaB helicase structure^15^ (**Fig. 2h**). However, the seam (identified by the unoccupied nucleotide-binding site) in the gp41 C-tier is at the interface between the A-B and E-F dimers, whereas the seam is within the E-F dimer in the DnaB helicase. Interestingly, the active gp41 helicase is more planar, with a vertical offset between the highest and lowest subunits of only 5 Å, as compared to a 20 Å offset between equivalent subunits in the DnaB helicase (**Supplementary Fig. 7**).

Each gp41 helicase CTD can coordinate two ssDNA backbone phosphates with sidechains Asn327–Tyr329 of the L1 loop, Ala372–Ala375 of the L2 loop, and Lys358 (**Fig. 3a**). Overall, the ssDNA coils around the interior of the gp41 CTDs coordinated by the L1 and L2 loops in a similar manner in both the spiral and planar forms of the gp41 helicase; however, the ssDNA becomes more coiled as the helicase switches from the open spiral to the planar ring (**Fig. 3b**). In the open spiral gp41 helicase, the ssDNA binds to five (subunits B-F) of the six helicase subunits and coils inside the right-handed C-tier spiral, as compared to the activated helicase ring in which the sixth subunit (subunit A) now joins the other five subunits to coordinate the ssDNA phosphate backbone (**Fig. 3c-e, Supplementary Fig. 8**). Therefore, the 10 nt of the ssDNA stabilized in the inactive open spiral is increased to a total of 12 nt stabilized in the active closed ring gp41 helicase. The diameter of the DNA coil is ∼23 Å, similar to the dimensions of the ssDNA observed in the T7 gp4, the bacterial DnaB, and the yeast CMG helicases (**Fig. 3f**). Therefore, the gp41 helicase likely unwinds DNA with a mechanism similar to the T7 and DnaB helicases^20,24^.

**Figure 3.**
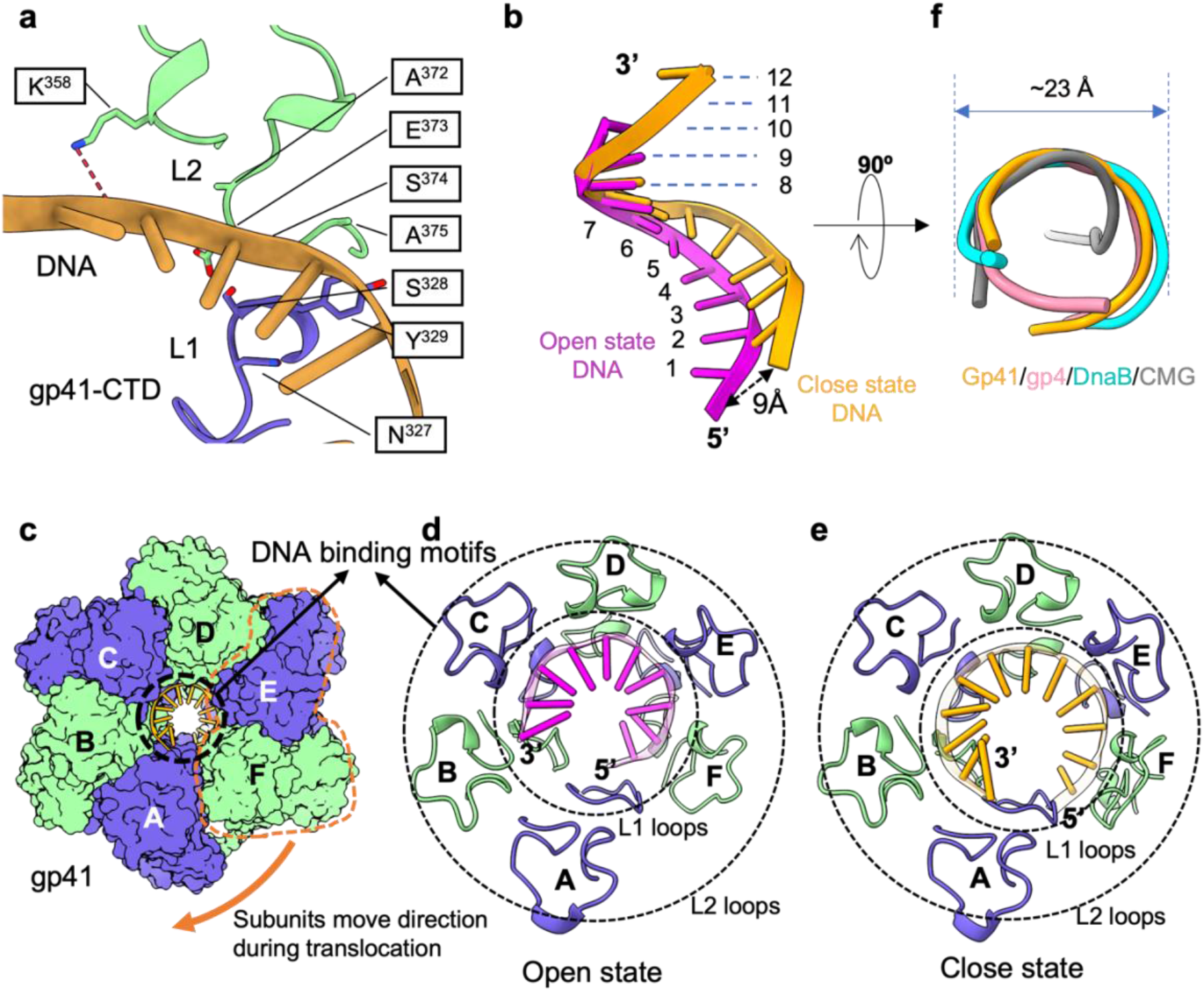
Comparison of ssDNA binding in the inactive open spiral and active closed-ring states of the gp41 helicase. **a)**The ssDNA backbone is sandwiched in the gp41 CTD between the tip of the α-helix following L1 loop (blue) and L2 loop (green). **b)** Superimposition of the ssDNA in the two states demonstrates that the ssDNA end moves by as much as 9 Å to adopt a flatter conformation in the closed-ring state. **c)** Top view of the gp41 helicase showing the coiled ssDNA in the central channel stabilized by the gp41 CTDs. The ssDNA interacts with five gp41 CTDs in the open state (**d**), and with six gp41 CTDs in the closed state (**e**). The DNA interacting loops are between the two concentric dotted circles and shown in cartoons. The DNA bases are shown in cartoon and in sticks. **f**) Superimposition of the ssDNA in the T4 helicase with ssDNA inside the T7 gp4 (PDB entry 6N9V), bacterial DnaB (PDB entry 4ESV), and yeast CMG helicase (PDB entry 5U8T). These ssDNA adopt a B-DNA-like configuration with a similar diameter of ∼23 Å.

### Bipartite interaction between the primase and the helicase hexamer

The final step of primosome assembly is for the gp61 primase to bind to the gp41 helicase hexamer. Importantly, the scissor-like movement that activates the gp41 helicase exposes a hydrophobic patch on the side of the gp41 helical hairpins, which recruits the gp61 primase (**Supplementary Fig. 9**). Therefore, the gp41 helicase in the active closed-ring form, but not the inactive open-spiral form, can recruit a primase. The primosome map 3-II shows the gp61 primase bound to the active, planar gp41 helicase hexamer. Interestingly, the structure of the gp41 helicase in the primosome complex is highly similar to the closed-ring helicase observed in the absence of gp61 primase (map 2-II), suggesting that the primase binding to the activated closed-ring gp41 helicase does not cause additional major conformational changes.

By classifying the primosome particles based on the gp61 primase regions and refining the entire primosome particles in each class, we obtained three primosome EM maps (**Fig. 4, Supplementary Figs. 4 and 5**). In all three maps, the gp41 helicase has a higher local resolution of 2.7 Å than the overall primosome resolution of 3.3 Å, 3.2 Å, and 3.5 Å, respectively. This indicates the partial flexibility of the primase subunits, especially at the distal RPD region; however, sidechain level details are present at the gp41 helicase/gp61 primase interface (**Supplementary Fig. 10a-b**). Aided by AlphaFold2, we were able to build atomic models of a T4 primosome in the three EM maps with a resolved full-length primase. The three primosome structures are similar, but not identical, to each other because of the asymmetry of the seam present in the gp41 helicase; they differ in the location of the two gp41 helicase dimers that bind the gp61 primase (**Fig. 4**). The gp61 ZBD and RPD bind the gp41 helicase similarly in the three binding modes with the ZBD binding to the first N-terminal helical hairpin dimer and the RPD binding to the second helical hairpin dimer in a clockwise direction when viewed from the primase side of the primosome. The three spatial and stationary binding modes of gp61 primase may represent three temporal steps of a primosome as the gp41 helicase translocates on ssDNA.

**Figure 4.**
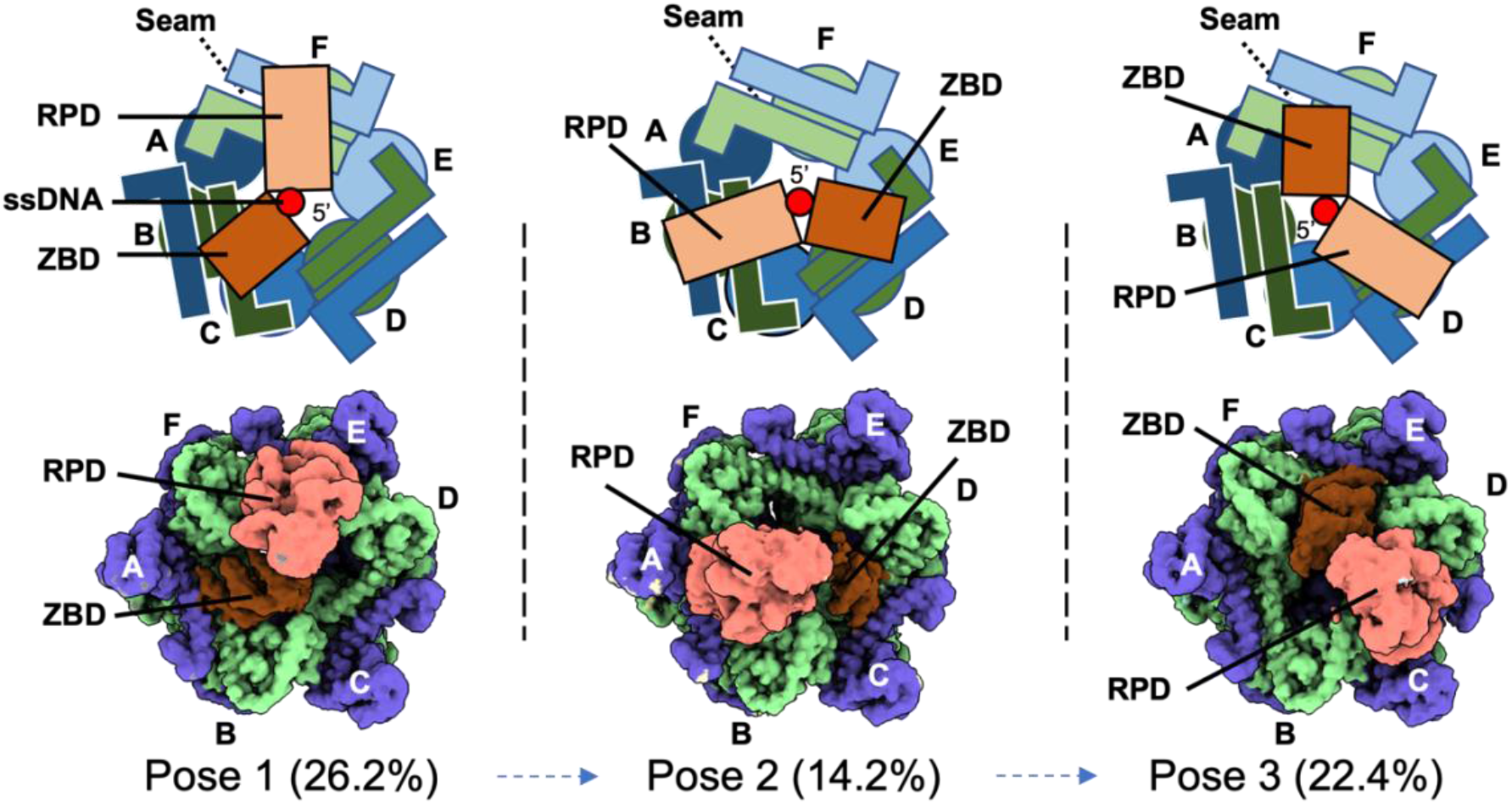
Three gp61 primase binding poses on the gp41 helicase. The three gp61 primase binding poses are observed in the T4 primosome in the presence of ssDNA and ATPγS. The upper panels are sketches of the three EM maps shown in the lower panels. One gp61 primase has two binding sites on the helicase hexamer: the gp61 ZBD binds to two NTDs of one gp41 dimer, and the gp61 RPD binds to two NTDs of a second gp41 dimer. The two NTDs of the third gp41 dimer are unoccupied. In each binding pose, the ssDNA 5′-end emerging from the gp41 helicase consistently passes between the gp61 ZBD and RPD domains. The particle population of each pose is given in parentheses. These poses are similar, but not identical, and are referred to as binding poses 1 through 3. The poses are non-equivalent because of the asymmetric helicase structure due to the presence of a seam between subunits A and F.

We found that ∼16% of the primosome particles had gp61 primase flexibly associated with it such that the gp61 primase density/location could not be accurately determined. In addition, a subpopulation of primosome particles (4.9%) was identified with only the gp61 RPD stably bound to the gp41 helicase, while the position of the gp61 ZBD was too disordered to be seen and was presumed to be flexible in solution (**Supplementary Fig. 4**). The stoichiometry of the T4 primosome has been controversial, with reports ranging from 6:1^30,37^ to 6:6^31,38^ helicase to primase subunits in the complex. The bipartite binding mode of gp61 primase to gp41 helicase observed in the majority of the primosome particles is consistent with a stoichiometry of 6 helicase:1 primase. But static structures can underestimated the complexity of protein quaternary structures and/or miss the dynamic equilibrium between different quaternary forms of complexes in solution^39^. The observation of subpopulations of primosome particles with other than bipartite or undeterminable primase binding modes to helicase suggests that other than 6:1 or flexible stoichiometries are also possible for the T4 primosome in solution.

In our structure, the gp61 primase interacts with the gp41 helicase in a bipartite manner with both the ZBD and the RPD (**Fig. 5a-b**). The gp61 ZBD interacts with the gp41 helicase via a helicase-interacting motif (HIM1; Ile74–Lys97) consisting of a helix-turn-helix motif following the Zn-ribbon core, and the RPD interacts with the gp41 helicase via a second helicase-interacting motif (HIM2; Ala327–Lys341) consisting of a single α-helix at the C-terminus following the catalytic TOPRIM fold (**Fig. 5a**). The gp61 HIM1 and HIM2 motifs bind to a similar region on separate gp41 NTD dimers (**Fig. 5b**). The gp61 HIM1 binds primarily to the gp41 helical hairpin, but also contacts the globular subdomain of the gp41 NTD (**Fig. 5c**). Specifically, Lys79, Glu80, Phe81, and Pro83 in the short first α-helix and the turn region of the gp61 HIM1 form Van der Waals interactions with the gp41 globular subdomain; and the longer second α-helix of the gp61 HIM1 forms an extensive interface with the gp41 helical hairpin, involving the Tyr86-Arg94 region of gp61 primase and the Phe104–Glu118 region of gp41 helicase. Sequence alignment reveals that the gp61 HIM1 is not conserved, suggesting that the interaction between the primase ZBD and the helicase may be unique to the T4 primosome (**Supplementary Fig. 11**). The gp61 HIM2 forms a helix-helix packing interaction with the N-terminal helical hairpin of the gp41 NTD dimer (**Fig. 5d**). The interface is largely hydrophobic, but also includes two hydrogen bonds - one between Lys333 of gp61 and Ser108 of gp41 and the other between Ser337 of gp61 and Thr115 of gp41. Comparison of the T4 gp61 primase and the bacterial DnaG primase reveals that the gp61 primase contains the first two domains of the DnaG primase but lacks the third domain of the bacterial primase (**Supplementary Fig. 12a**). The third DnaG primase domain functions to interact with the DnaB helicase and the ssDNA-binding protein^40^. Therefore, the interaction between the gp61 primase and gp41 helicase is expected to be different from the bacterial system. In contrast to the single-helix gp61 HIM2, the bacterial DnaG primase uses a C-terminal four-helix bundle as the HID to interact with DnaB^19^ (**Supplementary Fig. 12b-c**).

**Figure 5.**
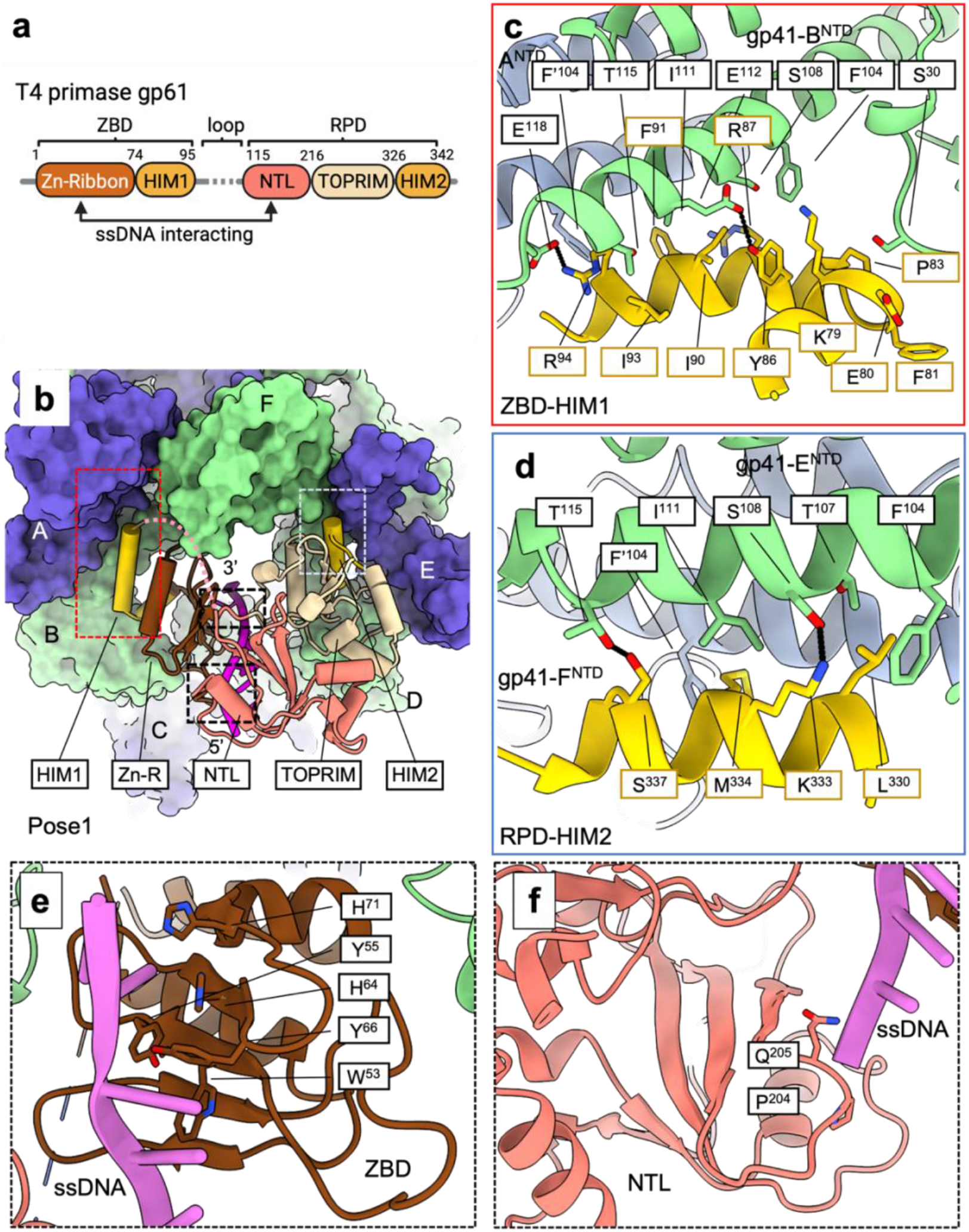
Interaction of the gp61 primase with the active and closed-ring gp41 helicase. **a)** A detailed architecture of the gp61 primase indicating the NTL: N-terminal lobe of the RPD and the TOPRIM, a signature domain conserved in all primases. **b**) A tilted bottom view of the T4 primosome showing the gp61 primase in cartoon binding to the NTDs of the gp41 helicase in surface. The binding mode is essentially the same in all three gp61 binding poses. The gp61 domains are colored as in (a). Enlarged views showing the detailed interactions between gp41 NTDs and **c)** the HIM1 and **d)** the HIM2 of the gp61 primase. The interacting residue pairs are connected by dashed black lines. Enlarged views showing detailed interactions between ssDNA and **e)** the zinc-ribbon core and **f)** the NTL in the RPD of the gp61 primase.

In our initial primosome structure, there was no density for the RNA primer, but the density for the ssDNA was present stretching from the gp61 ZBD to the N-terminal lobe (NTL) of the gp61 RPD (**Figs. 5b**); therefore, this structure may represent an ssDNA-scanning configuration of the primosome in search of a priming recognition sequence. The ZBD zinc-ribbon core is highly conserved and recognizes primer start sites on the template DNA^22,41,42^. In our structure, several residues with bulky side chains (Trp53, Tyr55, His64, Tyr66, and His71) of the zinc-ribbon core interact with the bases of the ssDNA, similar to the interactions between the T7 gp4 ZBD and ssDNA (**Fig. 5e**, **Supplementary Fig. 10c-d**)^24^. In the gp61 RPD, the topoisomerase-primase fold (TOPRIM; Thr217–Ile326) is sandwiched between the α/β-folded NTL (Lys115 to Ala216) and the HIM2 motif (Ala-327–Ile-342) (**Fig. 5a**). TOPRIM is the catalytic site for phosphate transfer and is conserved among the bacterial and phage primases. We found that the gp61 NTL binds the ssDNA via the edge of the β-sheet core, similar to the ssDNA binding by the bacterial DnaG primase^20^; however, the bacterial DnaG NTL is longer and contains a second DNA binding site, a β-hairpin motif, that is absent in the gp61 RPD. Nevertheless, the ssDNA in the DnaG RPD would be an extension of the ssDNA emerging from the gp41 helicase in our primosome structure, indicating a similar ssDNA-binding mode for the bacterial and phage primases (**Fig. 5f, Supplementary Fig. 10e-f**).

### Possible primer length determinant

We hypothesized that the absence of the RNA primer density in our primosome EM map was due to slow ATPγS hydrolysis by the WT gp41 helicase. Therefore, we utilized an inactive gp41 helicase mutant with an E227Q substitution (**Supplementary Fig. 13**) in the walker-B motif that abolished the ATP-hydrolysis activity^24^, and assembled a T4 primosome with the WT gp61 primase and the ssDNA/RNA primer substrate. We separately refined the helicase and primase regions to improve the resolution (**Supplementary Figs. 14-15**). The mutant gp41 helicase region had a resolution of 3.6 Å and revealed a structure essentially the same as the WT helicase (**Supplementary Fig. 14c**). The gp61 primase region achieved ∼4.0 Å local resolution where the pentaribonucleotide primer was observed to form a short hybrid duplex with the template DNA that was absent in the WT primosome map (**Fig. 6a-b**). The RNA primer density was just enough to accommodate five ribonucleotides, consistent with the RNA primer provided. Interestingly, we found that the gp61 RPD rotated around the HIM2 by ∼6° away from the ZBD to engage the ssDNA/RNA duplex, as compared to the gp61 primase in the WT primosome where the RNA primer is missing (**Fig. 6a-b**). Superposition of the gp61 primase structures of the WT and mutant primosomes shows that the RPD moves outward by 4 Å to accommodate the ssDNA/RNA duplex in the mutant structure (**Fig. 6c**).

**Figure 6.**
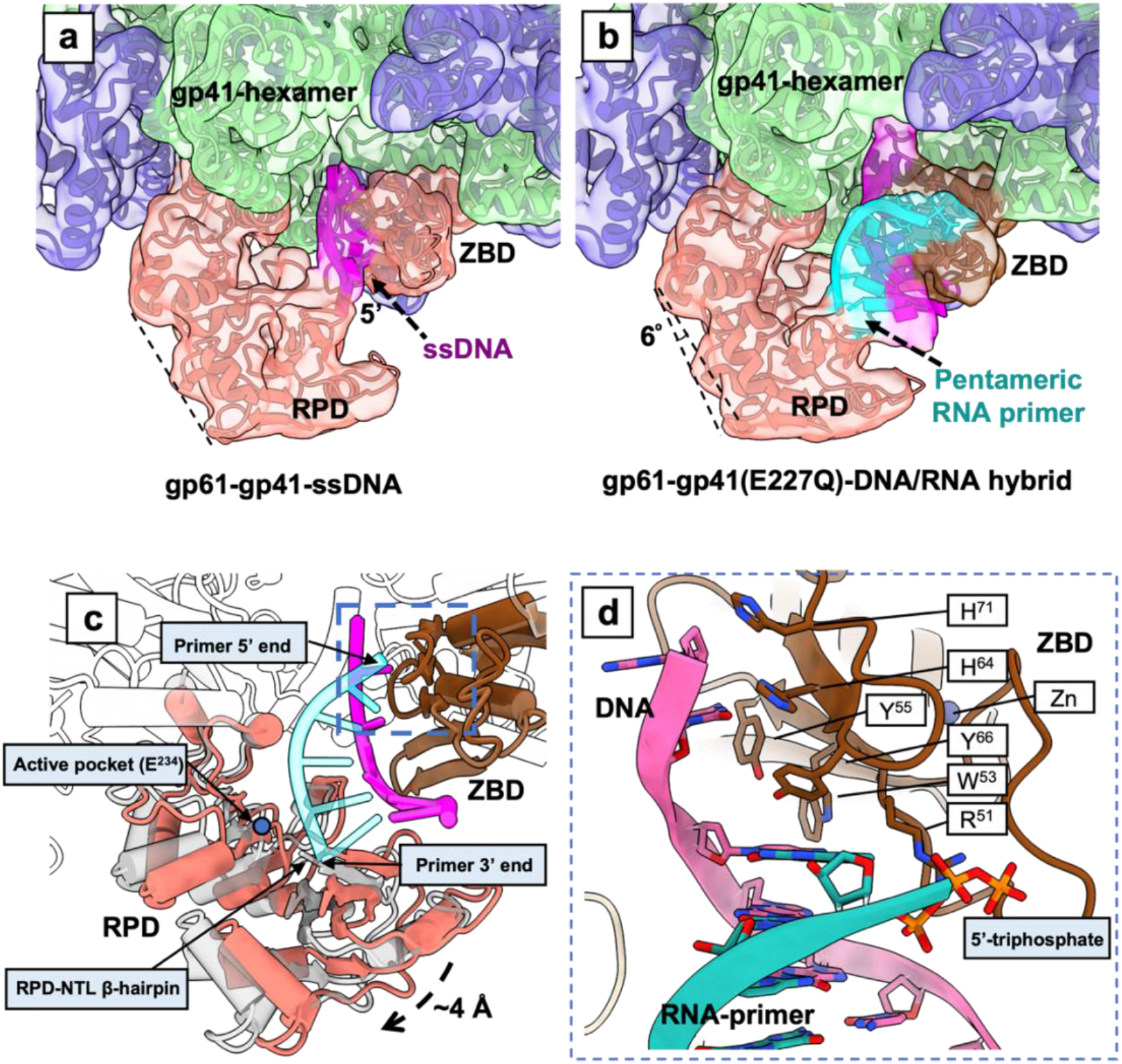
Two conformations of the T4 primosome captured by cryo-EM. Comparison of EM maps and models of **a)** the WT primosome with a resolved ssDNA and **b)** the helicase mutant (E227Q) primosome with a resolved ssDNA/RNA primer duplex. The EM maps are shown in transparent while the structural models are shown in cartoon. **c)** Superimposition of the two primosome structures showing that the gp61 ZBD is stationary, while the gp61 RPD rotates outwards by 4 Å to accommodate the 5-nt RNA primer. The structures are aligned by the gp41 NTD. Note that the 3′-end of the primer has passed the primase catalytic site − the E234 that is marked by a blue dot. The structures are colored as in (b) except that gp61 RPD of the mutant primosome is shown in transparent grey. **d)** Enlarged view of the dashed blue box region in (c) showing the interactions between the gp61 ZBD and the ssDNA/RNA primer. All labeled residues interact with the DNA (magenta) except for Arg51 and the preceding loop that interact with the 5′-triphosphate group of the RNA primer (green).

In the mutant primosome, rotation of the gp61 RPD caused the ssDNA template to rotate and interact more extensively with the gp61 ZBD as compared to the ssDNA in the WT primosome. The gp61 ZBD residues Trp53, Tyr55, His64, Tyr66, and His71 form multiple stacking interactions with the DNA bases (**Fig. 6c-d**). The 5-nt RNA primer extends from the gp61 ZBD to the NTL of the gp61 RPD. The primer 5’-triphosphate group is stabilized by Arg51 in the gp61 ZBD loop region (**Fig. 6d**); similar to the interaction observed in the T7 gp4 ZBD–ssDNA/RNA primer structure^24^. Therefore, the ZBD-mediated coordination of the 5′-end of the RNA primer is likely conserved among the phage priming systems. In the gp61 RPD, the RNA primer binds to the NTL and displaces the ssDNA bound there in the WT primosome structure. The 3’-end of the primer is stabilized by the β-hairpin of the gp61 RPD (**Fig. 6c**). Interestingly, the 3′-OH of the primer has passed over the catalytic site in the mutant structure, such that no additional ribonucleotides can be added in this configuration. This is consistent with the predominant 5-nt length of primers synthesized by the gp61 primase. We therefore assigned the mutant T4 primosome structure as in a post primer-synthesis state.

A working T4 primosome moves in a 5′ to 3′ direction on the lagging-strand DNA while simultaneously synthesizing an RNA primer in the opposing 3′ to 5′ direction. The gp61 RPD is physically attached to the gp41 helicase via its HIM2 and – as we have described above – can only rotate to accommodate a growing RNA primer. However, the gp61 RPD is connected by a linker loop (Lys97 to Lys115) to the stationary gp61 ZBD docked on the helicase. As the gp61 RPD rotates to synthesize and follow the elongating 3’-end of the primer, the linker loop becomes stretched, placing a limit on the range that the domain can follow the primer 3′-end. Indeed, we found that the linker loop stretched and became ordered in the mutant T4 primosome bound to a completed 5-nt RNA primer, as compared to the disordered linker loop in the ssDNA template-bound WT primosome (**Supplementary Fig. 16**). Therefore, the linker loop appears to contribute to the “counting” or “measuring” function that defines the primer length of the T4 primosome.

To test this hypothesis, a series of mutant primases was designed with either shortened or extended linker loops between the ZBD and RPD that included both conservative (e.g., the deletion or insertion of 1-2 residues) and substantial (e.g., the deletion or insertion of 4-5 residues) changes to the linker length, as well as the insertion of flexible (e.g., GGGGS sequence) or rigid (e.g., a repeat of the current linker sequence of PKEL) residues into the linker (**Supplemental Table 2**). The RNA primer synthesized by the WT primase and the 5 linker loop mutants was predominantly a pentamer (93.9 ± 0.9%) with very minor amounts of hexamer (2.8 ± 0.3%) or tetramer (3.3 ± 0.7%) RNA primers observed (**Supplementary Fig. 17**). Only in the case of mutant linker 2, where the linker loop was shortened by four residues, was an appreciable effect on the length of the primer noticed. While the predominant RNA primer was still a pentamer (95.2 ± 0.9%), the amount of hexamer primer (1.2 ± 0.1%) decreased and the amount of tetramer primer (3.7 ± 0.8%) increased with respect to the WT and other mutant primases as one would expect if the shorter linker loop restricted the rotation of the RPD thereby limiting the length of the primer that could be synthesized.

## DISCUSSION

Our systematic structure analysis has provided insights into the assembly process of the T4 primosome and its simultaneous DNA translocating and RNA priming mechanism. Based on the intermediate structures described above, we propose a multi-step T4 primosome assembly process (**Fig. 7a, Supplementary Video 1**). The assembly begins with the ATP-dependent oligomerization of gp41 helicase monomers. The helicase first assembles a right-handed hexameric open spiral with a lateral gap of 10 Å at its narrowest region. This gap likely serves as the lateral gate for the lagging-strand DNA to pass through. Existence of an open spiral structure is consistent with an earlier observation that the T4 helicase, but not the DnaB helicase, can self-load onto a naked ssDNA, although it does so inefficiently^43^ and therefore pairs with the gp59 helicase loader for efficient DNA loading in vivo. Interestingly, ssDNA binding in the helicase central channel does not immediately induce gate closure, suggesting that the ssDNA-bound open spiral is a metastable state and that there is an energy barrier between the open spiral and the gate-closed, active ring configuration. However, transitioning from the spiral to the planar ring does not appear to require ATP hydrolysis, because only ATPγS and not ADP molecules are present in both the spiral and ring forms. Therefore, we suggest that the binding energy of the N-tier helical hairpin dimers post the “scissor-like” motion drives the inactive-to-active conformational transition. And the binding energy of the two additional DNA nucleotides inside the active helicase further stabilizes the active configuration. This binding-energy-driven transition hypothesis is supported by our observation that the mutant gp41 helicase, lacking ATP hydrolytic activity, also adopted the active planar ring in the assembled primosome. Importantly, the conformational change in the N-tier helical hairpin dimers exposes an otherwise hidden hydrophobic surface to recruit gp61 primase to assemble the primosome. It is possible that the cryptic primase-binding site in the helicase may have evolved to prevent the primase from binding until after the helicase is activated and encircles an ssDNA. Docking of a gp61 primase to the helicase hexamer completes the assembly of a functional primosome.

**Figure 7.**
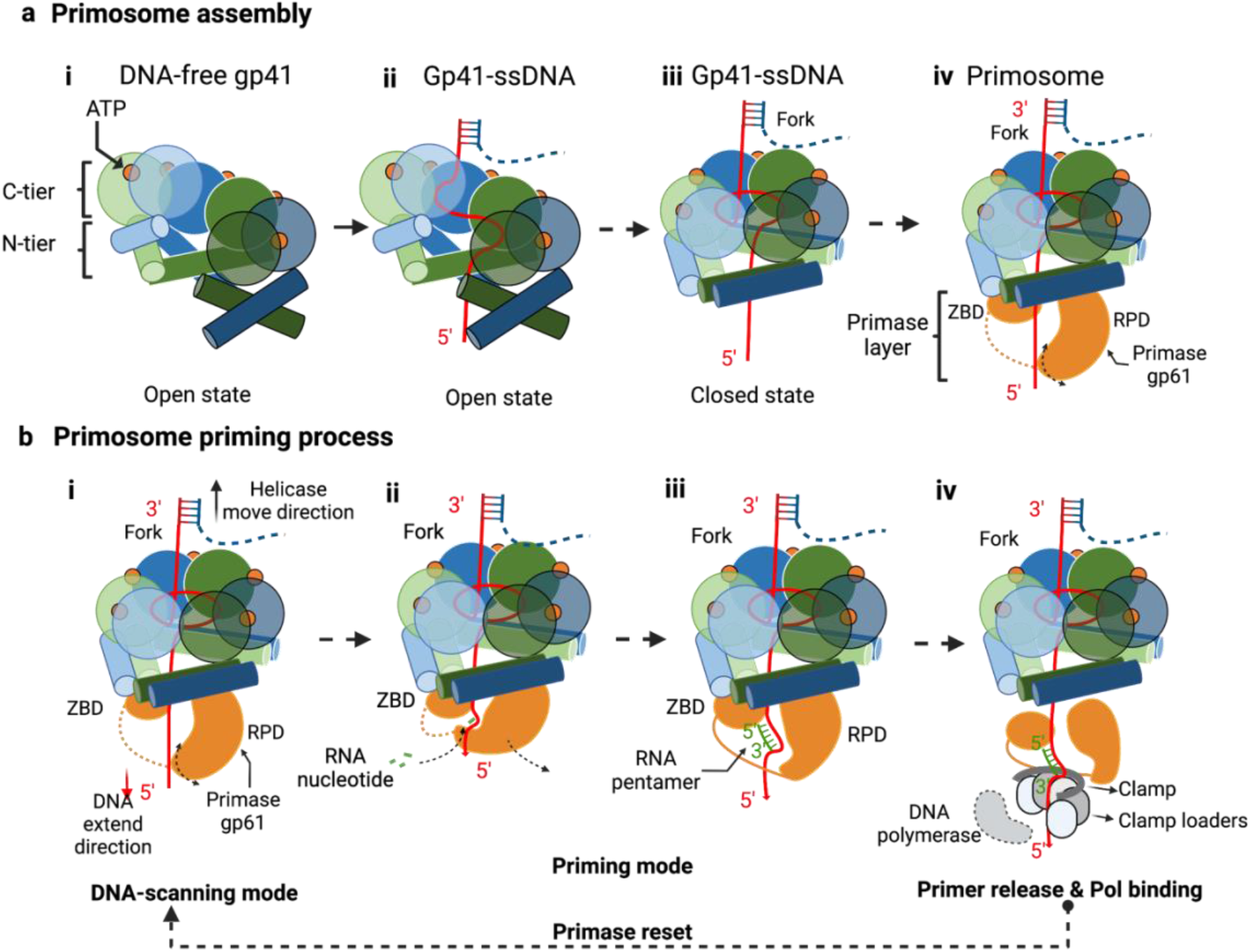
Hypothetical model for primosome assembly and DNA-unwinding coupled RNA priming of the T4 replication system. **a)**The four-step primosome assembly process is based on the four observed intermediate structures in this study. Note that the helicase loader gp59 is omitted in the scheme but is required in vivo. **b)** A proposed four-step DNA-unwinding coupled RNA priming mechanism (steps i to iv); see text for details.

We have described the T4 primosome in two distinct states (**Supplementary Video 2**). The first state is observed in the WT primosome in which the primase interacts with the ssDNA template in three largely equivalent poses. We suggest this is a DNA-scanning mode of the primosome in which the primase ZBD scans the ssDNA sequence for a 5’-GTT or 5’-GCT primer start site^29^ (**Fig. 7b**, step i). Once a primer start site is detected, we expect that the gp61 catalytic RPD will swing towards the ZBD where the primer start site is bound and initiate primer synthesis (**Fig. 7b**, step ii). This step is hypothetical as we did not experimentally capture this state in our cryo-EM grids. Because DNA translocation by the gp41 helicase (5’ to 3’) and primer synthesis by the gp61 primase (3’ to 5’) occur in opposite directions on the lagging-strand DNA, the gp61 RPD will need to rotate away from the ZBD during primer synthesis to maintain contact with the nascent 3’-end of the RNA primer. Indeed, after the 5-nt RNA primer (in the RNA/DNA hybrid duplex) has been fully synthesized, the RPD has rotated to the furthest possible extent away from the ZBD such that the linker loop connecting the two domains appears to be stretched and becomes ordered in the cryo-EM map (**Fig. 7b**, step iii). We propose that the linker loop contributes to the “measuring tape” function of the gp61 primase that limits the primer to 5 nt in the T4 system. Shortening the linker loop by four residues indeed affected the length of the synthesized primer. However, the effect was subtle; slightly less hexameric primer and more tetrameric primer was synthesized, albeit the pentameric primer was still the predominant product. Therefore, other factors besides the linker loop likely influence the length of the primer synthesized by the T4 primase. One potential factor is the affinity of the primase to the helicase. We suggest that a lengthening primer/template may exert an increasing force on the primase. Upon reaching the full length, the primase is pushed away and dissociates from the helicase, leading to the termination of primer synthesis. Such a scenario is consistent with the knowledge that T4 primase alone has little activity and is fully active only in complex with the helicase in the context of the primosome ^44,45^.

At the end of the priming reaction when a 5-nt RNA primer has been synthesized, the RNA primer/ssDNA must be handed over to the lagging-strand DNA polymerase through a mechanism that is currently unknown. Previous studies have found that handoff to the lagging-strand polymerase is stochastic and is successful between 20% and 40% of the time depending on the gp45 clamp and gp44/62 clamp loader levels^46^. We have also shown that a fraction of RNA primer/gp61primase complex dissociates from the gp41 helicase suggesting that the helicase/primase affinity in vitro is in the micromolar range^32^. This dissociated primer/primase complex remains on the lagging-strand replication loop and can act to block further replication of the lagging strand by the polymerase producing gaps in the replicated DNA for the T4 replisome^47^. We speculate that the post RNA primer-synthesis configuration with the completed RNA primer rotated past the primase active site allows the gp44/62 clamp loader to recognize and bind the 3’-OH to initiate primer handoff by loading the gp45 clamp and gp43 DNA polymerase on to the RNA primer/ssDNA (**Fig. 7b**, step iv). Alternatively, the elongating ssDNA emerging from the gp41 helicase applies pressure on the completed RNA primer/primase complex to dissociate from the helicase thereby creating a signal for lagging-strand polymerase recycling. A balance between RNA primer handoff to the polymerase and RNA primer/primase signals is necessary for coordinated leading- and lagging-strand synthesis in the T4 system. Further comment on the nature of the primer handoff will require cryo-EM analysis of the T4 replisome comprised of the helicase, primase, and DNA polymerase anchored by clamp on DNA.

## ACKNOWLEDGMENTS

Cryo-EM images were collected at the David Van Andel Advanced Cryo-Electron Microscopy Suite in the Van Andel Research Institute. We thank Gongpu Zhao and Xing Meng for facilitating the cryo-EM data collection. This work was supported by the US National Institutes of Health grants GM013306 (to S.J.B.) and GM131754 (to H.L.), and the Van Andel Institute (to H.L.).

## AUTHOR CONTRIBUTIONS

M.M.S., S.J.B., and H.L. conceived and designed experiments. X.F., M.M.S., and R.L.S performed experiments. All authors analyzed the data and participated in the manuscript preparation.

## DECLARATION OF INTERESTS

The authors declare no competing interests.

## DATA AND MATERIALS AVAILABILITY

The protein data bank accession codes for the atomic coordinates reported in this paper are 8DUO for the open spiral of the gp41 helicase hexamer; 8DTP for the closed ring of the gp41 helicase hexamer bound to ssDNA; 8DUE for the open spiral of the gp41 helicase hexamer bound to ssDNA; 8DVF, 8DVI, and 8DW6 for the T4 primosome in pose 1, pose 2, and pose 3, respectively; and 8G0Z, 8DWJ, and 8GAO for the helicase region, primase region and the whole primosome, respectively, of a mutant T4 primosome bound to an RNA primer/DNA hybrid in a post RNA primer-synthesis state. The EM data bank accession codes for the 3D EM maps reported in this paper are EMD-27724 for the open spiral of the gp41 helicase hexamer (map 1-I); EMD-27720 and EMD-27719 for the open spiral of the gp41 helicase hexamer bound to ssDNA (map 2-I and map 3-I); EMD-27708 and EMD-27707 for the closed ring of the gp41 helicase hexamer bound to ssDNA (map 2-II and map 3-II helicase region); and EMD-27737, EMD-27739, and EMD-27751 for the WT T4 primosome bound to ssDNA in poses 1, 2, and 3, respectively. EMDB-27707, EMDB-29744, and EMD-29902 are deposited for the helicase region, primase region and a composite map of the mutant T4 primosome bound to an RNA primer/DNA hybrid, respectively. EMDB-29744 is a local refinement from map EMDB-27756, which is deposited as well.

## METHOD

### Mutagenesis, Expression, and Purification of the T4 Proteins

The construction of expression plasmids for wt-gp59 helicase loader^48^, wt-gp41helicase^49^ and wt-gp61primase^31^ were described previously. The inactive gp41 helicase was generated by introducing the site-directed mutant E227Q into the gp41-intein fusion pET-IMPACT vector^49^. The mutation was made using the QuikChange Site-Directed Mutagenesis Kit (Agilent) with the mutagenic forward primer 5′-GTT CTT TAC ATT TCC ATG **<u>C</u>**AA ATG GCA GAA GAA GTC TG-3′ (the boldface underlined letter indicates the mutation site) and its reverse complement.

The series of gp61 primase linker loop mutants were generated by introducing insertions or deletions between Proline 106 and Lysine 111 of the linker loop into the gp61-intein fusion pET-IMPACT vector^31^. The modifications were made in the middle of the linker to avoid disrupting any interactions between the ZBD or RPD and the ends of the linker. The mutations were made using the QuikChange Multi Site-Directed Mutagenesis Kit (Agilent) with the mutagenic primers in supplementary table 2 (the underlined portions align to the WT primase sequence; the red dashes or bases indicate deletions or insertions, respectively).

**Supplementary Table 2.**
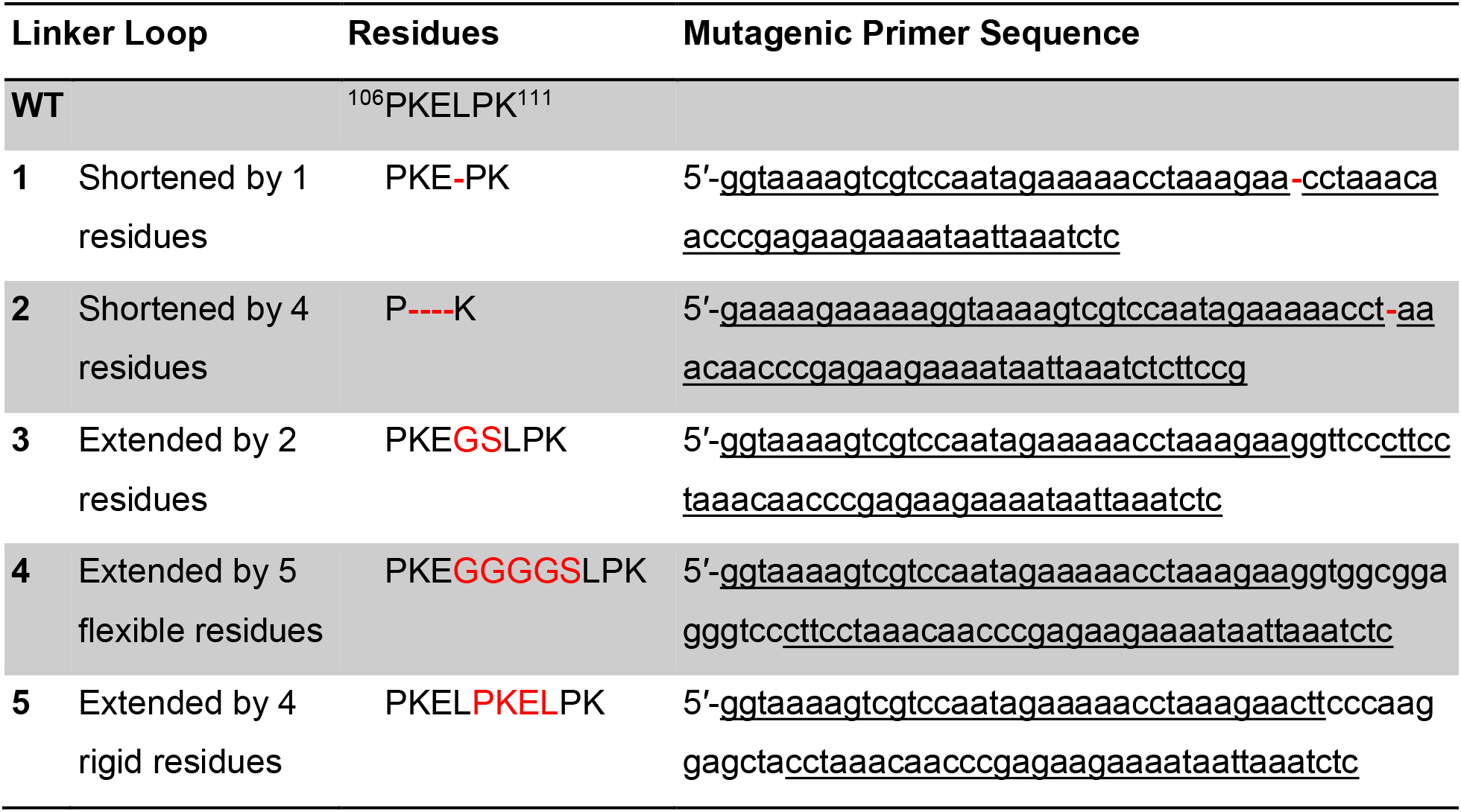
Table of gp61 primase linker loop mutants.

The self-cleaving, intein-based expression plasmids were transformed into *E. coli* BL21(DE3) cells and grown in NZCYM media at 37 °C to an optical density of 0.4 at 600 nm. The cultures were then cooled to 18 °C, and protein expression was induced with 0.4 mM isopropyl 1-thio-β-D-galactopyranoside. After 16 – 20 h of shaking, cells were harvested by centrifugation and either resuspended in chitin column binding/high salt buffer (20 mM TrisOAc pH 7.8, 1 M NaOAc, 0.1 mM EDTA, and 10% glycerol) with a cocktail of protease inhibitors (Roche) or frozen for protein purification later. Cells were lysed using sonication or high-pressure homogenization, and cell debris was pelleted at 40 000 × g. Cell-free extract was loaded onto chitin resin (New England Biolabs) for chitin-based affinity chromatography, and the chitin resin was washed with chitin column binding buffer. The resin was resuspended in cleavage/low salt buffer (20 mM TrisOAc pH 7.8, 100 mM NaOAc, 0.1 mM EDTA, and 10% glycerol) with 75 mM β-mercaptoethanol and incubated overnight at 4 °C to facilitate intein-mediated cleavage. The protein was eluted from the chitin column in low salt buffer for anion exchange (gp41 helicase) or cation exchange (gp59 helicase loader and gp61 primase) chromatography developed with a linear gradient of high salt buffer. The eluted protein was dialyzed into storage buffer (10 mM TrisOAc pH 7.8, 25 mM KOAc, 5 mM Mg(OAc)_2_, 2 mM dithiothreitol, and 20% glycerol) and analyzed for purity using SDS-PAGE. Protein concentrations were determined by measuring the absorbance at 280 nm using extinction coefficients based on the protein sequence. Proteins for cryo-EM were purified in buffers containing HEPES pH 7.8 instead of TrisOAc pH 7.8 and were frozen for storage immediately following ion exchange chromatography without dialysis into storage buffer.

### Protein Activity Assays

Helicase Unwinding Assay. The helicase unwinding assays were performed in replication buffer (25 mM TrisOAc pH 7.8, 150 mM KOAc, and 10 mM Mg(OAc)_2_) containing 50 nM unwinding fork DNA, 500 nM trap ssDNA, 2.5 mM ATP, 350 nM gp59 helicase loader, 300 nM gp61 primase, and various ratios of WT gp41: gp41(E227Q) totaling 300 nM helicase in a typical reaction volume of 60 μL. The reaction was carried out at 37 °C; 10 μL aliquots were withdrawn at 30 s time points over 2 min and quenched with an equal volume of loading buffer (240 mM EDTA, 0.2% SDS, 15% glycerol, 1 μg/mL bromophenol blue, and 1 μg/mL xylene cyanol FF). Reaction products were separated by 10% native PAGE in 1x TBE buffer and analyzed using a phosphorimager. The sequences of the oligonucleotide substrates were as follows: fork lead (5′-CAT CAT GCA GGA CAG TCG GAT CGC AGT CAG ATT TAC TGT GTC ATA TAG TAC GTG ATT CAG-3′); fork lag (5′-TAA CGT ATT CAA GAT ACC TCG TAC TCT GTA CTG ACT GCG ATC CGA CTG TCC TGC ATG ATG-3′); and trap (5′-CTG ACT GCG ATC CGA CTG TCC TGC ATG ATG-3′). The unwinding fork was made by mixing fork lead and fork lag DNA in equal molar amounts and radiolabeling the 5′-ends with T4 Polynucleotide Kinase and [γ-^32^P]ATP.

Primase Priming Assay. Priming reactions were performed in replication buffer containing 1.5 μM ssDNA oligo (71-mer), 4 mM ATP, 200 μM each CTP, GTP, and UTP, 8 μCi of [α-^32^P]CTP, 3.0 μM gp59 helicase loader, 3.0 μM gp41 helicase, and 1.5 μM WT or mutant gp61 primase in a typical reaction volume of 25 μL. The reactions were carried out at 37 °C; 5 μL aliquots were withdrawn at 1 min time points over 4 min and quenched with 15 μL loading buffer (167 mM EDTA, 67% formamide, and 1 μg/mL xylene cyanol FF). Priming products were separated by 20% denaturing urea-PAGE in 1x TBE buffer and analyzed using a phosphorimager. The sequence of the ssDNA oligonucleotide substrate was 5′-AGA GGG AGA TTT AGA TGA GAT GAT TGA GGA TGG AGA T<u>GT T</u>GA TGG AGA GAT GAT GAA TGA TGA GAT GAG GG-3′ and contained a GTT priming recognition site (underlined).

### DNA and RNA Constructs

All DNA oligos were purchased from Integrated DNA Technologies and either PAGE or HPLC purified. A ssDNA oligo 5′-GAA TGA GGA GTA GTA GTG AAT GTA GTG AGG TAA TAT CG<u>G CT</u>G GTC TGG TCT GTG CCA AGT TGC TGCAAA A-3′ containing a GCT priming recognition site (underlined) was used in cryo-EM. The corresponding RNA primer 5′-ppp-rGrCrCrGrA-3′ with a 5′-triphosphate moiety was synthesized using a standard run-off transcription protocol with the T7 RNA polymerase as previously described^47^. The ssDNA and RNA primer were annealed together by heating to 90 °C and slowly cooling to 4 °C at the final concentration of 250 μM.

### In vitro assembly of the T4 primosome

We assembled the T4 primosome step-by-step with purified components as previously described^30^. We first exchanged the stocks of gp41 helicase and gp61 primase to the in vitro reaction buffer (20 mM HEPES pH 7.8, 100 mM NaCl, 10 mM MgCl_2_ and 2 mM DTT) by centrifugal filtering with Amicon Ultra-0.5 devices (10 kDa cutoff). We started with the assembly of 6 μM gp41 helicase by adding 5 mM ATPγS (step 1). We next added 12 μM ssDNA/RNA primer substrate (step 2). In step 3, we added 3 μM gp61 primase to assemble the primosome. The mixtures were incubated for 30 min at room temperature between steps before the reaction products were withdrawn for cryo-EM grid preparation.

### Cryo-EM grid preparation and data collection

To alleviate any preferred orientation of the particles, 0.2% n-Octyl-β-D-glucoside was quickly mixed with the various primosome assembly samples before preparing the grids. Aliquots (4 μL) of the primosome assembly samples were applied to glow-discharged holey carbon grids (Quantifoil R2/1 Copper, 300 mesh) in the climate-controlled chamber of a FEI Vitrobot Mark IV. The EM grids were blotted for 3 s with filter paper and then plunged into liquid ethane and stored in liquid nitrogen. Pilot datasets of around 300 micrographs were collected on a 200-kV FEI Arctica electron microscope equipped with a K2 summit camera (Gatan) for screening purposes. The 3D reconstruction and refinement led to preliminary 3D maps with resolutions around 6 Å to confirm the quality of the grids. Two individual datasets for each primosome assembly sample were then collected using SerialEM^50^ on a TFS Titan Krios electron microscope operated at 300 kV and at a nominal magnification of 130,000× equipped with a K3 summit camera (Gatan) using the objective lens defocus range of –1.0 to –2.0 μm. All the EM images were recorded in the super-resolution counting and movie mode with a dose rate of 0.88 electrons per Å^2^ per frame; a total of 75 frames were recorded in each movie micrograph.

### Image processing

The gp41-helicase alone dataset contained 150 micrographs for screening purposes. A dataset of 50,003 particles was selected after 3D classification and 3D refined, and polished to the final map with 5.7 Å resolution.

The gp41 helicase-DNA/RNA primer and primosome datasets, containing 4091 and 11476 movie stacks, respectively, were processed in the same fashion. The movies were drift-corrected with electron-dose weighting and two-fold binned using MotionCor-2.1^51^. The full datasets were split into subsets with around 2000 movie stacks and imported into Relion-3^52^. In each dataset the particles were auto picked based on templates from the scanning results and extracted with 4-fold binning. The auto picked particles were then imported into Cryosparc2 (v3.2)^53^ for 2D classification. The “good” 2D class averages with defined structural features were selected as input for *ab initio* 3D model reconstruction. From this stage, the particles were automatically classified into open states and closed states for separate refinement. For each state, the particle images from the best 3D maps reconstructed from subset of the data were merged and converted to the RELION format using UCSF PyEM (https://github.com/asarnow/pyem). At this stage, the gp41 helicase-DNA/RNA primer dataset had 783,172 and 406,763 particles in the open and closed states, respectively. The gp41 helicase-gp61 primase-DNA/RNA primer dataset had 1,904,089 and 1,920,895 particles in the open and closed states, respectively. For each state, the re-extracted original-scale particles were filtered through another round of 3D classification using RELION3.1 before 3D refinement. Then the particles were further CTF-refined, polished, 3D refined again and postprocessed to reach the final density map. To resolve the gp61 primase density map, another round of heterogeneous classification using Cryosparc2 (v3.2)^53^ was run on the particles that belonged to the closed gp41 helicase in the gp41 helicase-gp61 primase-DNA/RNA primer datasets revealing one class of gp61 primase binding to gp41 helicase. This density map provided the mask to extract only the gp61 primase signal in all of the closed-state particles through RELION3.1. Then 3D classification without alignment was applied to the extracted partial particles revealing three major binding orientations of the gp61 primase (pose 1, 2 and 3) and a minor single gp61 RPD binding mode (pose 4). Lastly, the partial particles from each orientation were reverted to the full particles, followed by 3D classification to select the particles that give the best results. The final density maps were generated after another 3D refinement.

To estimate the number of particles with bound primase, we set the closed-ring helicase hexamer particles required for primase binding as 100%, which were classified into three classes; two of which had EM density for the primase (84% combined) and one class had no EM density for the primase (16%). From the 84% particles with a bound primase, we further separated these particles into three primase poses (1 – 3) (26.2%, 14.2% and 22.4%, respectively) and additional primase with density only for the RPD domain (4.9%). Combined, these classes accounted for 67.7% in the 84% of the particles with electron density for primase. Therefore, the remaining 16.3% of the particles must have primase flexibly bound to the helicase hexamer such that no specific orientation could be determined.

The mutant primosome was processed similarly. A total of 10,224 movies were collected. We first obtained a subset of 340,403 particle images with the helicase in the closed state, and then used the wild-type gp41 helicase-gp61 primase structure as a template to perform another round of 3D classification, leading to a mutant primosome EM map that reached an overall resolution of 3.9 Å. Close inspection of the map revealed that the template DNA density inside the mutant helicase was discontinuous, indicative of the presence of multiple helicase poses that likely resemble those observed in the wild-type helicase. We did not perform additional 3D classification because the dataset was already small, and our focus here was to capture the ssDNA/RNA primer bound to the primase. A focus refinement with Cryosparc2 (v.32)^53^ lead to a 4.1-Å resolution local map of primase/DNA-RNA hybrid. Then from the same pool of particles and using the gp41 helicase as a mask, we obtained an EM map for mutant gp41 helicase in the closed state at an overall resolution of 3.6 Å. This averaged structure is similar to the WT gp41 helicase. Both maps were postprocessed with DeepEMhancer^54^ with the *wideTarget* model and combined to generate a composite map using ChimeraX^55^.

The resolution of all final maps was calculated based on the 0.143 threshold of the gold standard Fourier shell correlation between the two independently constructed “half” maps, with each map using half of the dataset. The local resolution maps were calculated using RELION3.1 and displayed using UCSF ChimeraX.

### Model building and refinement

The high-resolution maps (better than 3.5Å) were postprocessed with DeepEMhancer with the *tightTarget* model for better density map details^54^. The initial atomic model for the gp41 helicase was built with comparative modeling while fitting into the electron density map with the Rosetta suite^56^ using the template of the DnaB helicase (PDB entry 6QEM). All the atomic models were then either manually rebuilt (like unfit regions, ligands and the ssDNA chain) or refined with the program COOT^57^ followed by real-space refinement in the PHENIX program^58^. The structures were further manually refined in COOT^57^ until no Ramachandran outliers could be identified. Finally, the atomic model was validated using MolProbity^59,60^. Because each state (open or closed) of the gp41 helicase in either the gp41 helicase-ssDNA or the gp41 helicase-gp61 primase-ssDNA datasets have similar conformations, the map with the best resolution for each state was used to build the model. The crossing angle between the gp41 NTD helical hairpins was calculated in Pymol (The PyMOL Molecular Graphics System, version 1.8.x; Schrödinger). The EM map of the ATPγS-bound, DNA-free gp41 helicase had a moderate resolution of 5.7 Å. The structural models of the open-state DNA-bound gp41 helicase subunits were then fit into the density map using UCSF ChimeraX^55^ and refined in the PHENIX program. The initial structural model of the gp61 primase was predicted with AlphaFold^61^. Due to the flexible linker loop hindering the prediction of the domain organization, the structural models of the gp61 ZBD and RPD were individually docked into the density map and refined with Phenix. The ssDNA/RNA primer binding conformation were modeled based on the structure of the T7 gp4 (PDB entry 6N9U) and refined within the density map. The structural model alignments and figures were generated using UCSF ChimeraX^55^, and the sketches were created using Biorender.com.

---

**Supplementary Table 1.** Cryo-EM data collection, refinement, and validation statistics.

|  | Close state gp41<br>hexamer with<br>ssDNA (Step 3)<br>(EMDB-27707)<br>(PDB 8DTP) | Close state gp41<br>hexamer with<br>ssDNA (Step 2)<br>(EMDB-27708) | Open state<br>gp41 hexamer<br>with ssDNA<br>(Step 3)<br>(EMDB-27719)<br>(PDB 8DUE) | Open state<br>gp41 hexamer<br>with ssDNA<br>(Step 2)<br>(EMDB-27720) |
| --- | --- | --- | --- | --- |
| **Data collection and processing** |  |  |  |  |
| Magnification | 105,000 | 105,000 | 105,000 | 105,000 |
| Voltage (kV) | 300 | 300 | 300 | 300 |
| Electron exposure (e<sup>-</sup>/Å<sup>2</sup>) | 66 | 66 | 66 | 66 |
| Defocus range (μm) | 1.0 to 2.0 | 1.0 to 2.0 | 1.0 to 2.0 | 1.0 to 2.0 |
| Pixel size (Å) | 0.828 | 0.828 | 0.828 | 0.828 |
| Symmetry imposed | C1 | C1 | C1 | C1 |
| Initial particle images (no.) | 4,802,236 | 1,314,307 | 4,802,236 | 1,314,307 |
| Final particle images (no.) | 848,518 | 233,875 | 690,193 | 268,443 |
| Map resolution (Å) | 2.7 | 3.4 | 2.9 | 3.5 |
| FSC threshold | 0.143 | 0.143 | 0.143 | 0.143 |
| Map resolution range (Å) | 2.0-5.0 | 2.0-7.0 | 2.0-7.0 | 2.0-7.0 |
| **Refinement** |  |  |  |  |
| Initial model used (PDB code) | 6QEM |  | 6QEM |  |
| Model resolution (Å) | 2.9 |  | 3.1 |  |
| FSC threshold | 0.5 |  | 0.5 |  |
| Model resolution range (Å) | 2.0-5.0 |  | 2.0-7.0 |  |
| Map sharpening *B* factor (Å<sup>2</sup>) | 83.0 |  | 73.9 |  |
| **Model composition** |  |  |  |  |
| Non-hydrogen atoms | 20944 |  | 20902 |  |
| Protein residues | 2592 |  | 2592 |  |
| Nucleotides | 12 |  | 10 |  |
| Ligands | MG: 5 AGS:5 |  | MG:5 AGS:5 |  |
| ***B* factors (Å<sup>2</sup>)** |  |  |  |  |
| Protein | 57.2 |  | 80.9 |  |
| Nucleotide | 95.8 |  | 69.1 |  |
| Ligand | 52.8 |  | 49.4 |  |
| **R.m.s. deviations** |  |  |  |  |
| Bond lengths (Å) | 0.004 |  | 0.003 |  |
| Bond angles (°) | 0.625 |  | 0.532 |  |
| **Validation** |  |  |  |  |
| MolProbity score | 2.06 |  | 2.42 |  |
| Clashscore | 9.28 |  | 12.72 |  |
| Poor rotamers (%) | 1.88 |  | 3.4 |  |
| **Ramachandran plot** |  |  |  |  |
| Favored (%) | 94.8 |  | 94.1 |  |
| Allowed (%) | 5.2 |  | 5.9 |  |
| Disallowed (%) | 0.0 |  | 0.0 |  |

|  | DNA-free gp41<br>hexamer<br>(EMDB-27724)<br>(PDB 8DUO) | Gp41-gp61-ssDNA (Pose 1)<br>(EMDB-27737)<br>(PDB 8DVF) | Gp41-gp61-ssDNA (Pose 2)<br>(EMDB-27739)<br>(PDB 8DVI) | Gp41-gp61-ssDNA (Pose 3)<br>(EMDB-27751)<br>(PDB 8DW6) |
| --- | --- | --- | --- | --- |
| **Data collection and processing** |  |  |  |  |
| Magnification | 36,000 | 105,000 |  |  |
| Voltage (kV) | 200 | 300 |  |  |
| Electron exposure (e<sup>-</sup>/Å<sup>2</sup>) | 48 | 66 |  |  |
| Defocus range (μm) | 1.4 to 2.4 | 1.0 to 2.0 |  |  |
| Pixel size (Å) | 1.16 | 0.828 |  |  |
| Symmetry imposed | C1 | C1 | C1 | C1 |
| Initial particle images (no.) | 71,335 | 4,802,236 |  |  |
| Final particle images (no.) | 50,003 | 124,592 | 97,947 | 142,027 |
| Map resolution (Å) | 5.7 | 3.3 | 3.2 | 3.5 |
| FSC threshold | 0.143 | 0.143 | 0.143 | 0.143 |
| Map resolution range (Å) | 4.0-10.0 | 3.0-7.0 | 3.0-7.0 | 3.0-7.0 |
| **Refinement** |  |  |  |  |
| Initial model used (PDB code) | 8DUE | 6QEM, 6N9U, 2AU3 | 6QEM, 6N9U, 2AU3 | 6QEM, 6N9U, 2AU3 |
| Model resolution (Å) |  |  |  |  |
| FSC threshold 0.5 | 7.7 | 3.6 | 3.5 | 4.1 |
| Model resolution range (Å) |  | 3.0-7.0 | 3.0-7.0 | 3.0-7.0 |
| Map sharpening *B* factor (Å<sup>2</sup>) | 210.0 | 99.4 | 90.7 | 109.0 |
| **Model composition** |  |  |  |  |
| Non-hydrogen atoms | 20704 | 23693 | 23693 | 23693 |
| Protein residues | 2592 | 2914 | 2914 | 2914 |
| Nucleotide | 0 | 17 | 17 | 17 |
| Ligands | MG: 5 AGS:5 | MG:5 AGS:5<br>ZN:1 | MG:5 AGS:5<br>ZN:1 | MG:5 AGS:5<br>ZN:1 |
| ***B* factors (Å<sup>2</sup>)** |  |  |  |  |
| Protein | 257.1 | 81.5 | 81.5 | 81.5 |
| Nucleotide | -- | 181.1 | 181.1 | 181.1 |
| Ligand | 205.6 | 52.7 | 52.7 | 52.7 |
| **R.m.s. deviations** |  |  |  |  |
| Bond lengths (Å) | 0.004 | 0.004 | 0.004 | 0.004 |
| Bond angles (°) | 0.737 | 0.666 | 0.667 | 0.622 |
| **Validation** |  |  |  |  |
| MolProbity score | 2.97 | 2.97 | 2.09 | 2.2 |
| Clashscore | 21.72 | 10.45 | 10.96 | 11.98 |
| Poor rotamers (%) | 7.1 | 1.78 | 1.78 | 3.2 |
| **Ramachandran plot** |  |  |  |  |
| Favored (%) | 91.9 | 95.1 | 95.1 | 95.1 |
| Allowed (%) | 8.1 | 4.9 | 4.9 | 4.9 |
| Disallowed (%) | 0.0 | 0.0 | 0.0 | 0.0 |
|  | Close state<br>mutated gp41<br>hexamer with<br>ssDNA<br>(EMDB-27707)<br>(PDB 8G0Z) | gp61-ssDNA/RNA<br>hybrids<br>(EMDB-27756)<br>(PDB 8DWJ) | Local refinement<br>of the primase<br>region in mutant<br>T4 primosome<br>(EMDB-29744) | Mutant<br>primosome with<br>E227Q helicase<br>composite map<br>(PDB 8GAO)<br>(EMD-29902) |
| **Data collection and processing** |  |  |  |  |
| Magnification | 130,000 | 130,000 | 130,000 |  |
| Voltage (kV) | 300 | 300 | 300 |  |
| Electron exposure (e-/Å<sup>2</sup>) | 80 | 80 | 80 |  |
| Defocus range (μm) | 1.0 to 2.0 | 1.0 to 2.0 | 1.0 to 2.0 |  |
| Pixel size (Å) | 1.029 | 1.029 | 1.029 |  |
| Symmetry imposed | C1 | C1 | C1 |  |
| Initial particle images<br>(no.) | 5,706,491 | 5,706,491 | 5,706,491 |  |
| Final particle images (no.) | 272,868 | 157,809 | 157,809 |  |
| Map resolution (Å) | 3.6 | 3.9 | 4.1 |  |
| FSC threshold | 0.143 | 0.143 | 0.143 |  |
| Map resolution range (Å) | 3.0-7.0 | 3.0-8.0 | 3.0-8.0 |  |
| **Refinement** |  |  |  |  |
| Initial model used (PDB<br>code) | 8DTP | 6N9U, 2AU3 | 6N9U, 2AU3 |  |
| Model resolution (Å) | 4.1 | 4.6 | 4.6 |  |
| FSC threshold | 0.5 | 0.5 | 0.5 |  |
| Model resolution range<br>(Å) | 3.0-7.0 | 3.0-7.0 | 3.0-8.0 |  |
| Map sharpening *B* factor<br>(Å<sup>2</sup>) | 121.6 | 126.3 | 70.5 |  |
| Model composition |  |  |  |  |
| Non-hydrogen atoms | 20944 | 3042 |  | 23987 |
| Protein residues | 2592 | 339 |  | 2931 |
| Nucleotides | 12 | 11 |  | 23 |
| Ligands | MG:5 AGS:5 | GTP:1<br>ZN:1 |  | MG: 5 AGS:5<br>ZN: 1 |
| *B* factors (Å<sup>2</sup>) |  |  |  |  |
| Protein | 61.1 | 243.1 |  | 82.8 |
| Nucleotide | 101.5 | 275.8 |  | 186.7 |
| Ligand | 57.1 | 270.3 |  | 93.6 |
| R.m.s. deviations |  |  |  |  |
| Bond lengths (Å) | 0.005 | 0.006 |  | 0.003 |
| Bond angles (°) | 0.721 | 1.268 |  | 0.592 |
| Validation |  |  |  |  |
| MolProbity score | 2.22 | 2.07 |  | 2.35 |
| Clashscore | 12.05 | 14.2 |  | 12.60 |
| Poor rotamers (%) | 2.1 | 0.33 |  | 3.47 |
| Ramachandran plot |  |  |  |  |
| Favored (%) | 94.4 | 93.77 |  | 94.55 |
| Allowed (%) | 5.6 | 5.64 |  | 5.45 |
| Disallowed (%) | 0.0 | 0.00 |  | 0.0 |

**Supplementary Figure 1.**
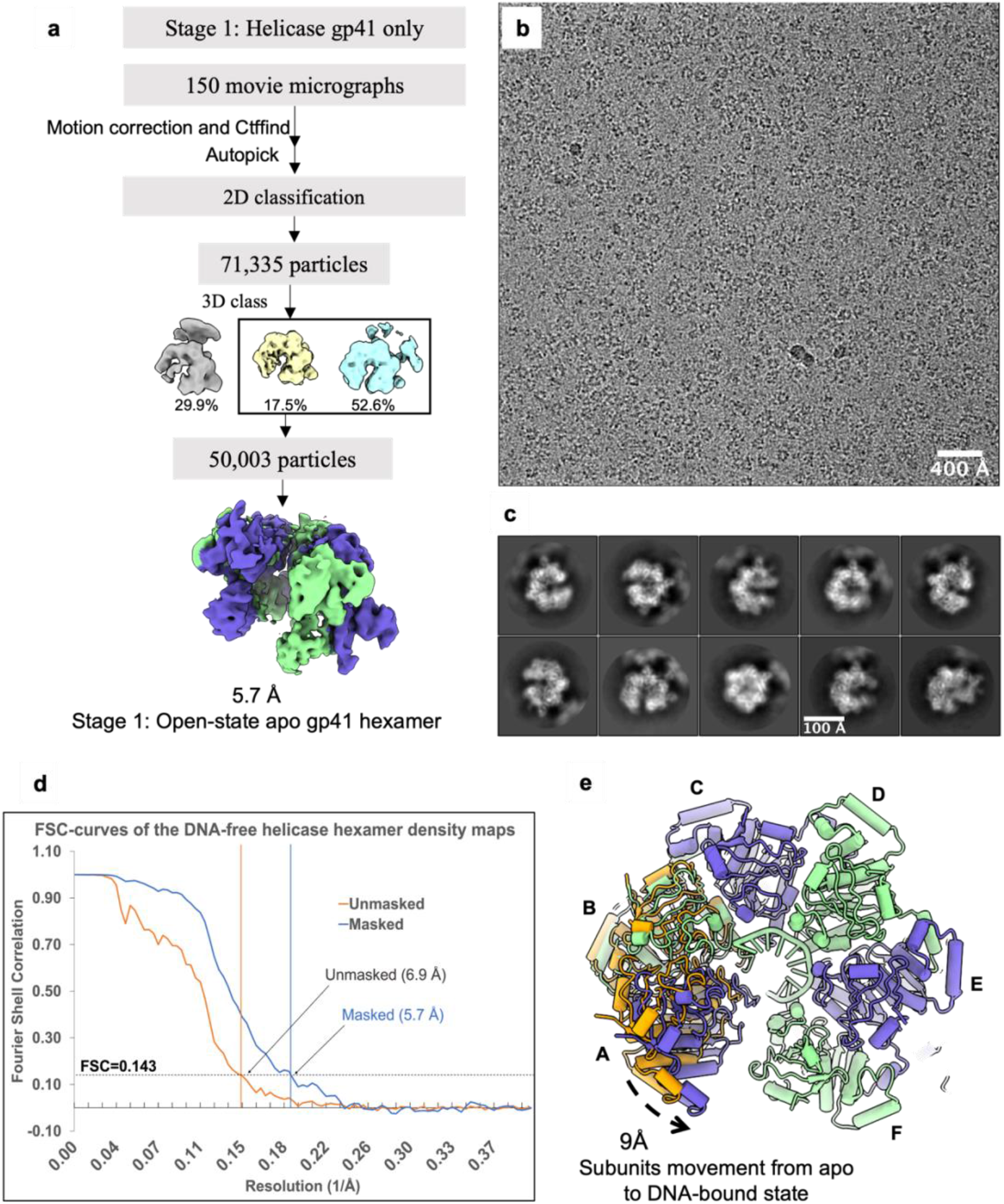
Processing of EM images of the ATPγS-bound gp41 helicase sample in the absence of ssDNA. **a)**Data processing flowchart. **b**) A typical raw micrograph. **c**) Selected 2D class averages. **d**) Gold-standard Fourier shell correlation curve indicating an averaged resolution of 5.7 Å at the 0.143 correlation threshold. **e**) Comparison of the open spiral gp41 helicase before and after binding ssDNA; the two structural models are aligned based on subunit F. Subunits A-F are colored alternatively in blue or green. Subunits A and B in the gp41 helicase structure without ssDNA are shown in orange to better show the movement of these subunits when binding to ssDNA.

**Supplementary Figure 2.**
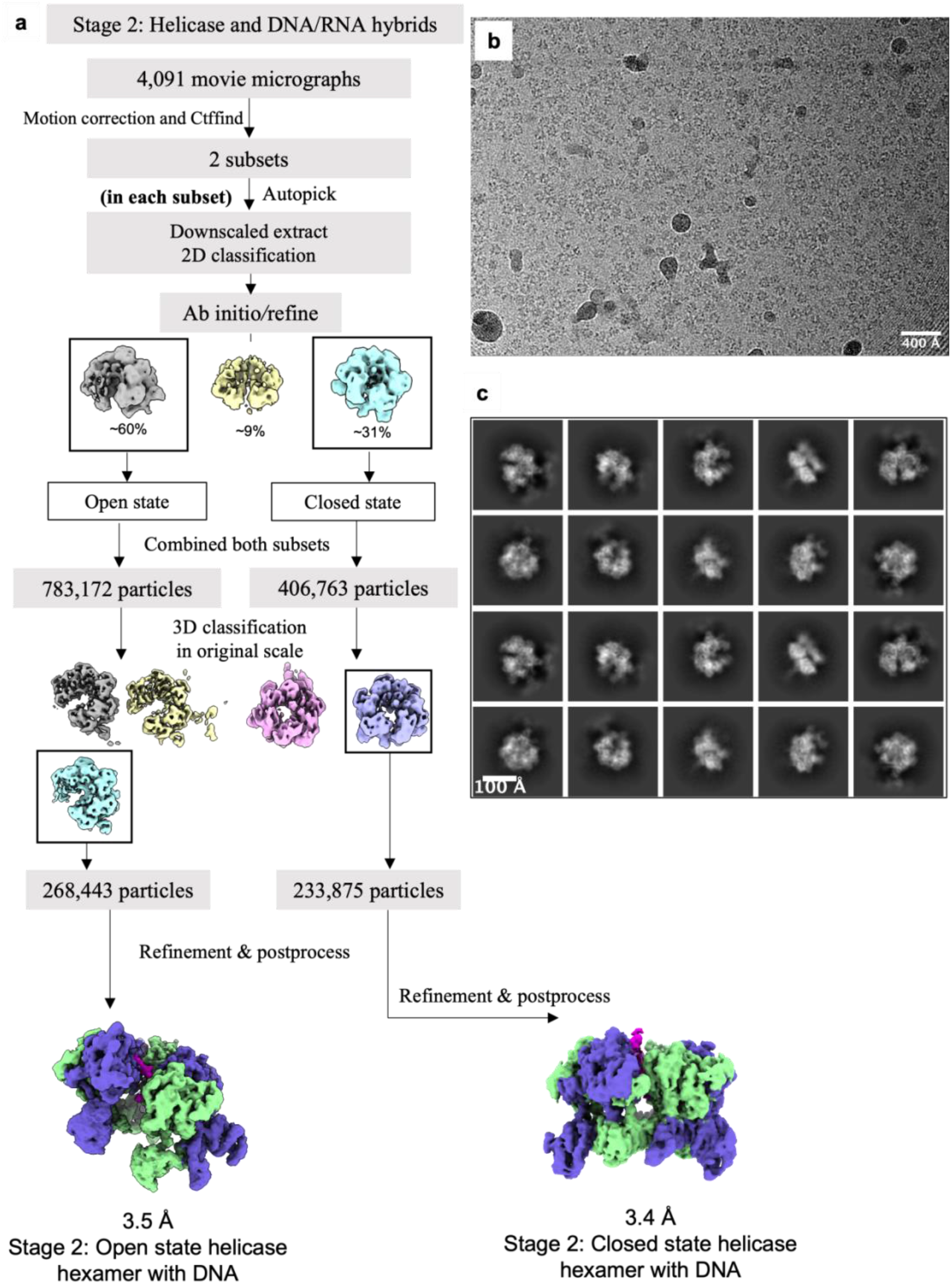
Processing of cryo-EM images from the sample of the gp41 helicase mixed with the ssDNA/RNA primer. **a)** Data processing flowchart. **b)** A typical raw micrograph. c) Selected 2D class averages.

**Supplementary Figure 3.**
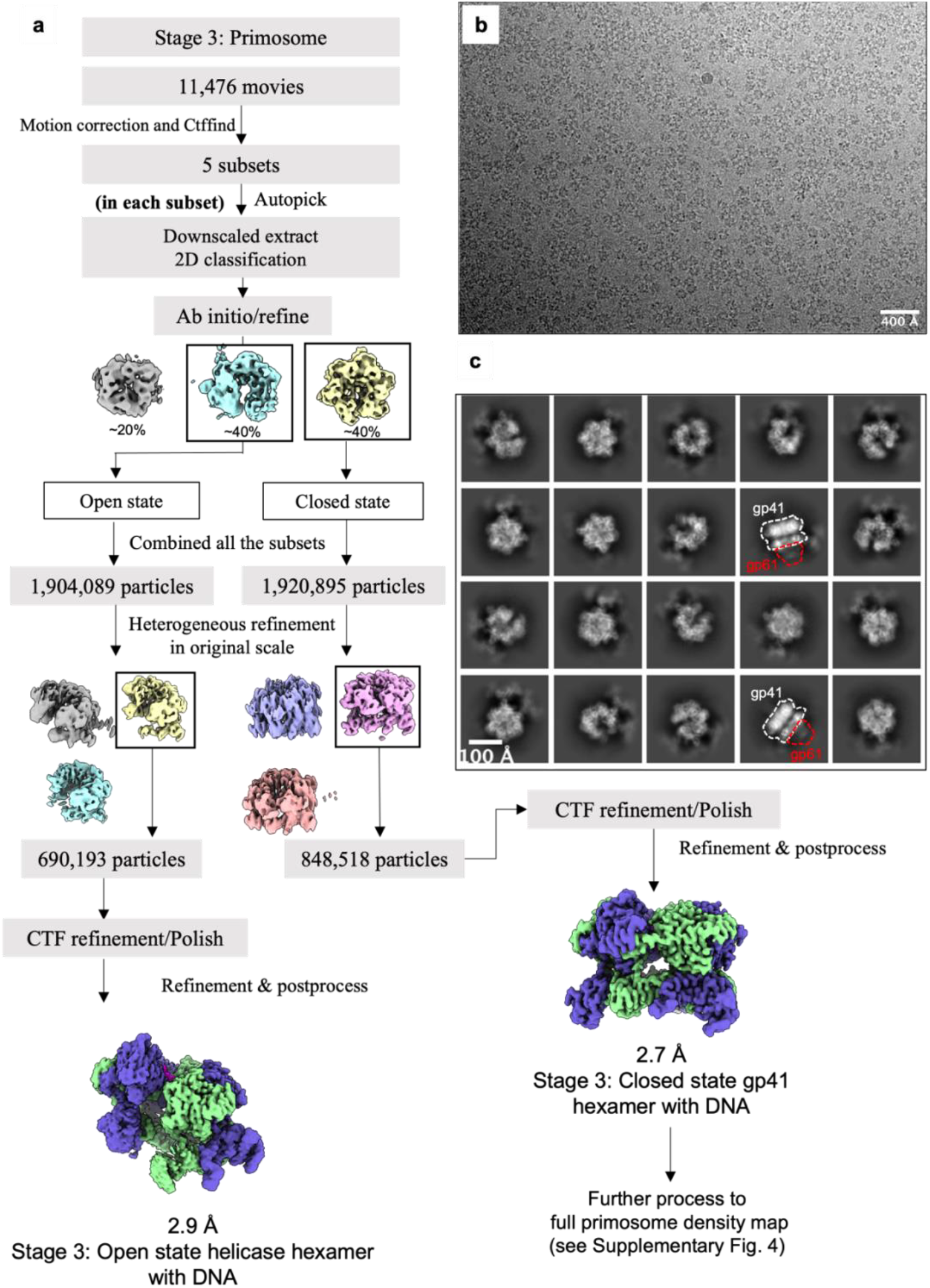
Initial processing of cryo-EM images from the primosome assembly sample. **a**) Data processing flowchart for reconstructing the gp41 helicase within the primosome particles. **b**) A typical raw micrograph. **c**) Selected 2D class averages showing the presence of the gp61 primase in some averages.

**Supplementary Figure 4.**
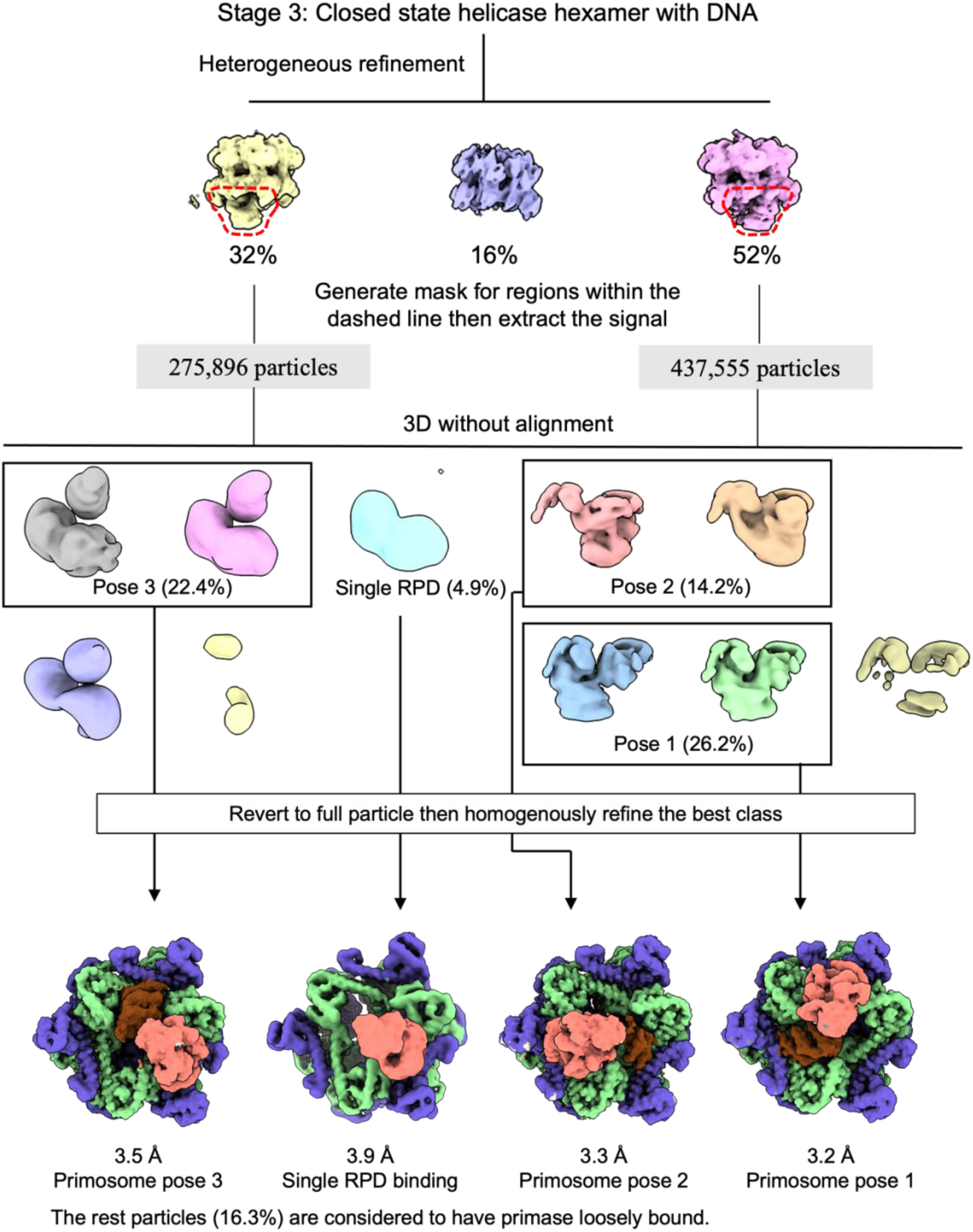
Additional processing of cryo-EM images from the primosome assembly sample. Further processing of the closed-state particles within the primosome sample led to the identification of three major bipartite binding poses and one minor single-domain (RPD) binding pose of the gp61 primase within the T4 primosome.

**Supplementary Figure 5.**
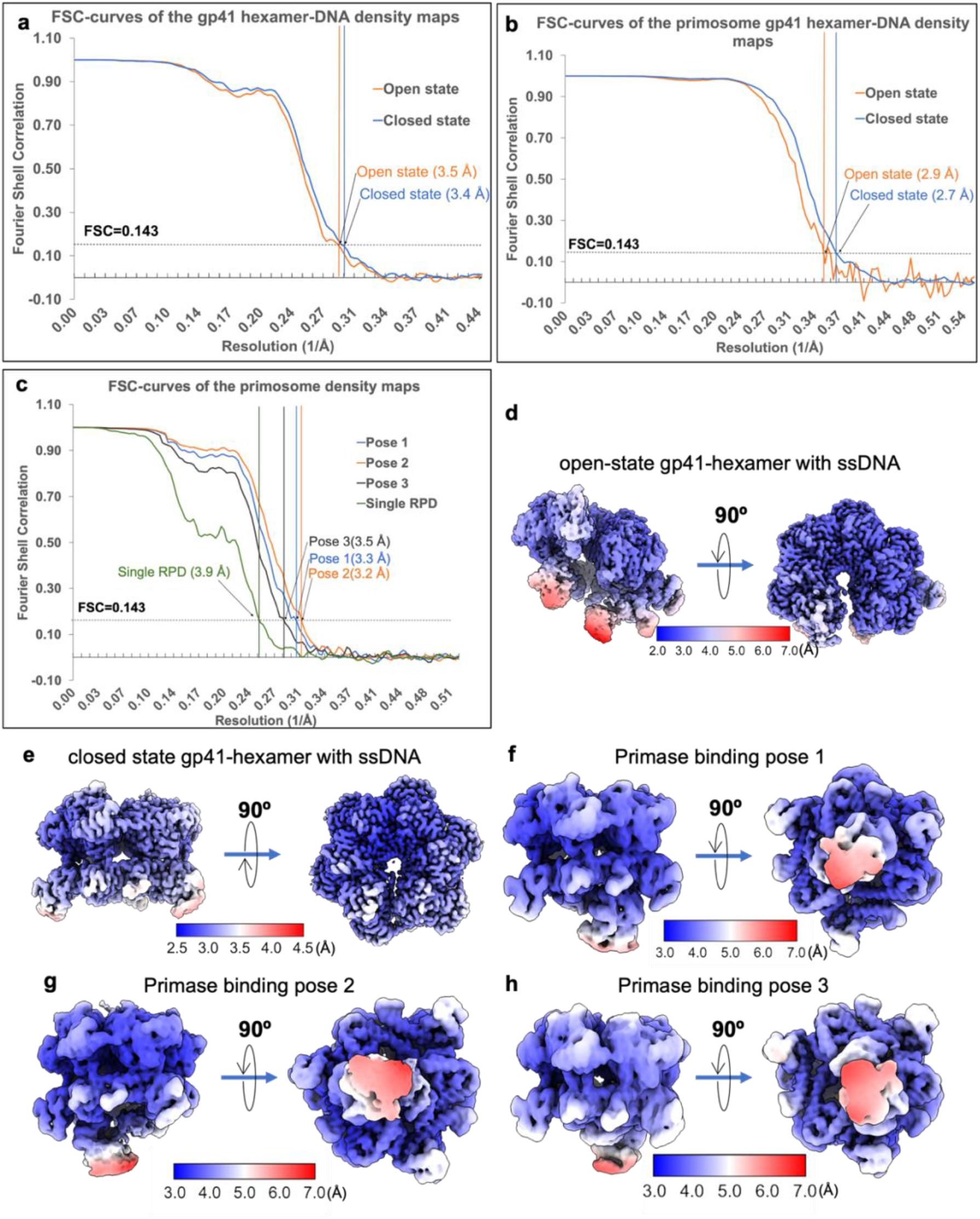
Resolution estimation of the EM maps of the gp41 helicase-ssDNA complex and the primosome. FSC curves of the open- and closed-state gp41 helicase/ssDNA in **a)** the gp41/ssDNA sample and **b)** the primosome sample. **c)** FSC curves of the four primase-bound poses in the primosome sample. Local resolution maps of the primosome assembly intermediates: **d)** the open-state and **e)** the closed-state of the gp41 helicase/ssDNA sample; and the primase-bound **f)** pose 1, **g)** pose 2, and **h)** pose 3 in the primosome sample.

**Supplementary Figure 6.**
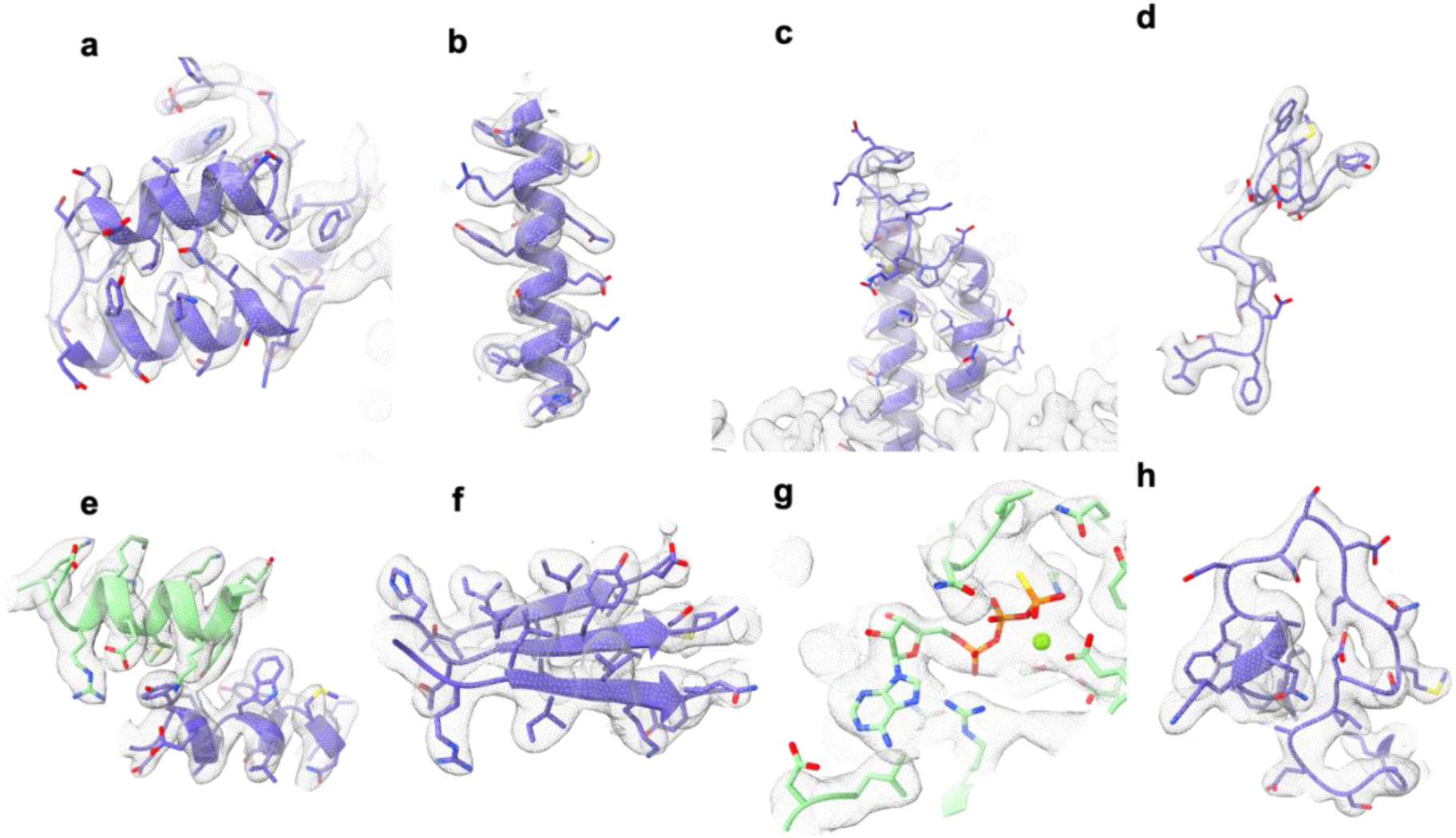
Fitting of the gp41 helicase atomic model with selected regions of the EM map. Comparison of the EM map rendered at a threshold of 2σ (semitransparent grey surface) with selected regions of the gp41 helicase atomic model: **a)** N-terminal globular subdomain; **b-c)** two regions from the N-terminal helical subdomain; **d)** the loop linking the NTD and the linking helix; **e)** the linking helix and the interacting helix of the adjacent subunit; **f)** the β-sheets within the RecA-like CTD; **g)** an ATPγS molecule and its binding pocket; and **h)** the DNA interacting loop 1. All density regions are extracted from the closed-ring helicase structure in EM map 3-II.

**Supplementary Figure 7.**
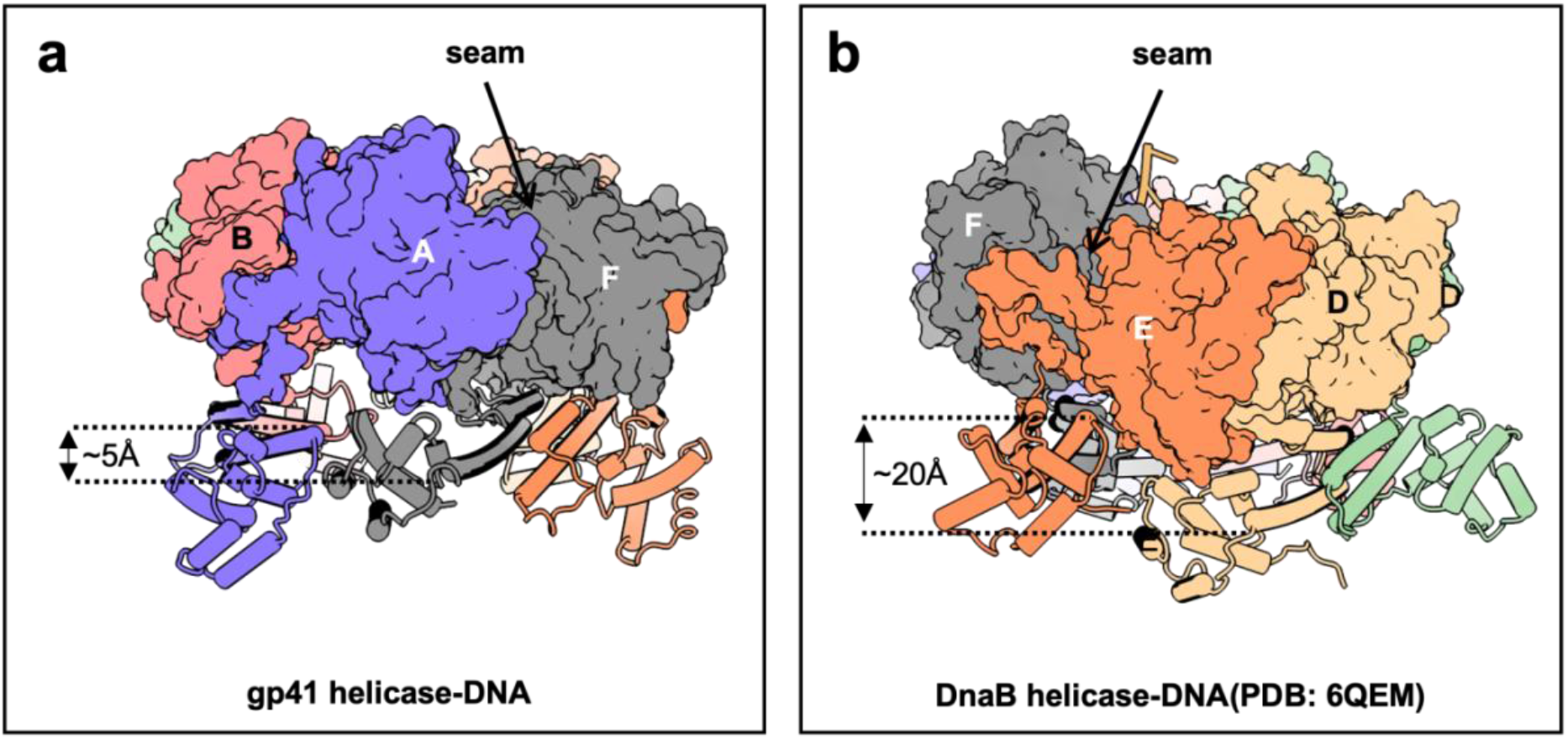
Comparison of the T4 gp41 and *E. coli* DnaB helicases in translocation-competent active states. Side views showing **a)** an almost planar ring arrangement of the gp41 helicase with a small (5 Å) lateral staggering (this study) and **b)** a twisted non-planar ring of the DnaB helicase with a large (20 Å) lateral staggering (PDB entry 6QEM) between the adjacent subunits. Both the gp41 and DnaB helicases assemble as trimer-of-dimers in their active state. The three dimers consist of subunit A-B, subunit C-D and subunit E-F. The seam identified as the nucleotide-free interface is between subunits A and F, which belong to two separate dimers, in the gp41 helicase and between subunits E and F, which belong to a single dimer, in the DnaB helicase.

**Supplementary Figure 8.**
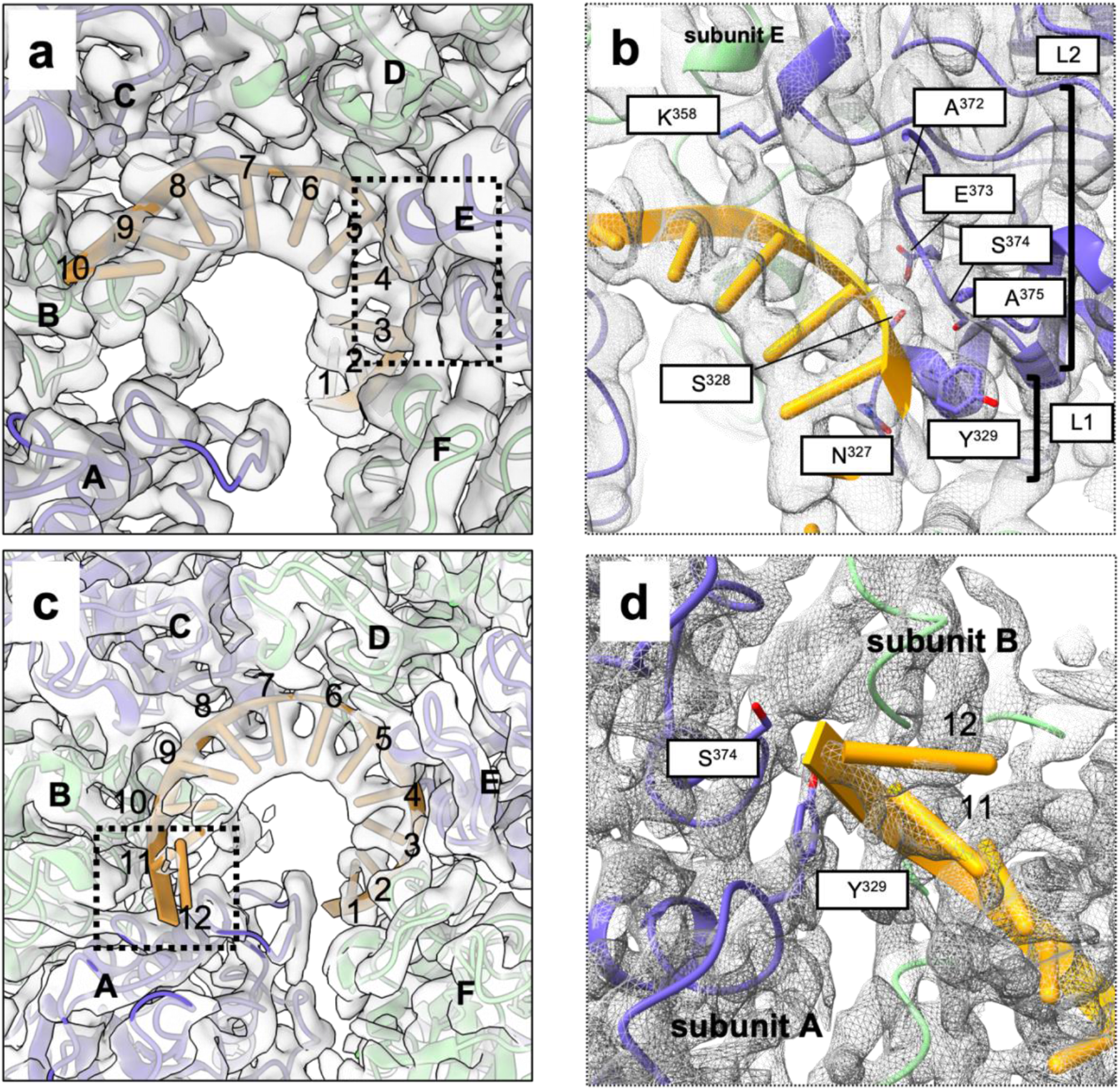
The ssDNA binding mode in the open and closed states of the gp41 helicase. **a)**An overall view and **b)** a zoomed-in view of the ssDNA binding region in the central channel of the open spiral gp41 helicase structure. **c)** An overall view and **d)** a zoomed-in view of the ssDNA binding region in the central chamber of the closed-ring gp41 helicase structure. The EM maps are shown as transparent grey surface in (a) and (c) and as grey meshes in (b) and (d); the atomic models of DNA are sticks; and the residues interacting with the DNA are sticks with the residue names labeled in one letter codes. L1 and L2 refer to the DNA-translocating loops 1 and 2, respectively.

**Supplementary Figure 9.**
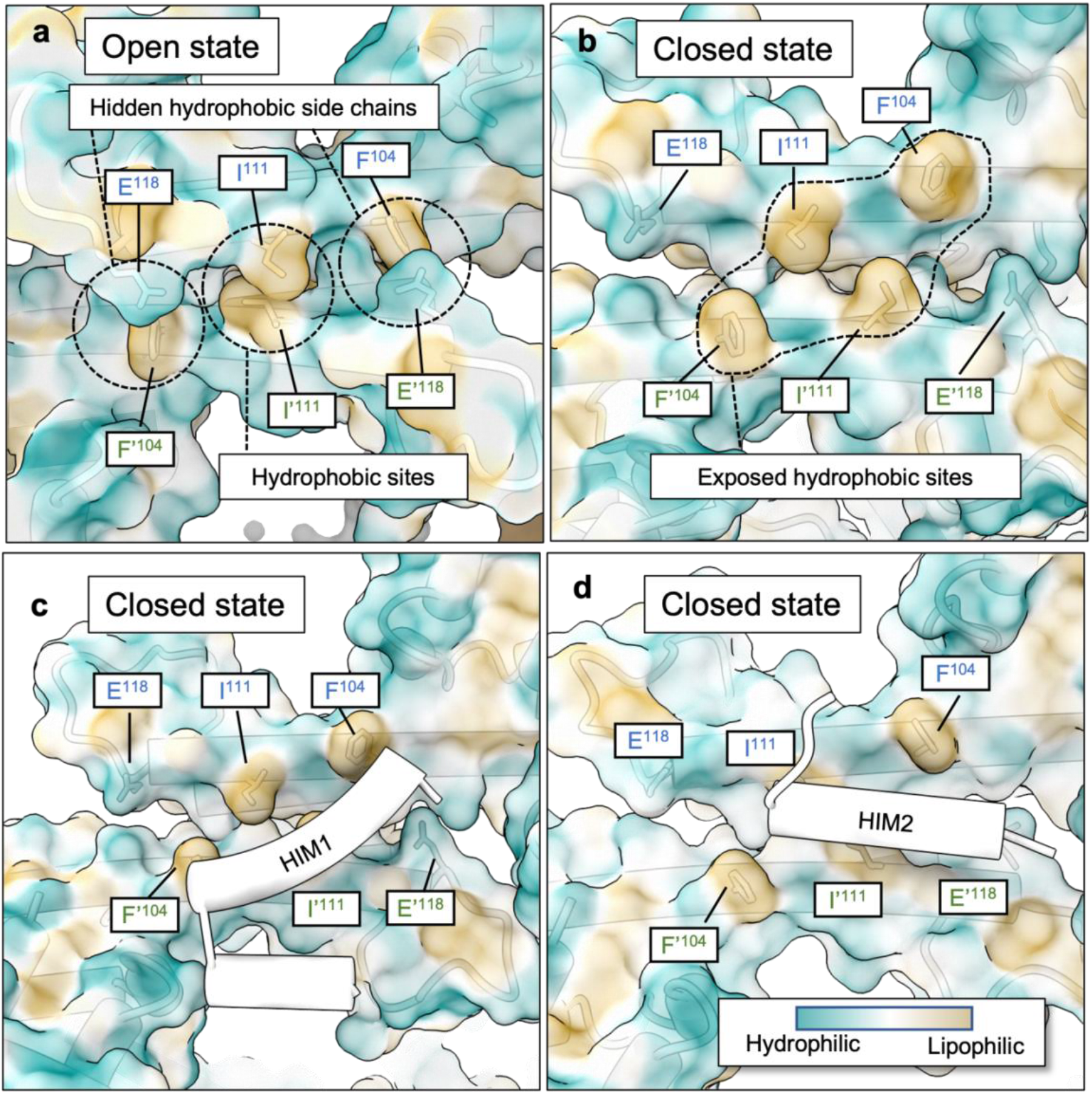
Surface hydrophobicity of the gp41 NTD dimers in various states. **a)**In the open spiral, the helical hairpins in one NTD dimer are in an “X” configuration where the two Ile111 residues from each subunit face each other, and the two hydrophobic Phe104 residues are shielded by the two hydrophilic Glu118 residues from each subunit. **b)** In the closed-ring gp41 helicase, the helical hairpins are parallel exposing the four hydrophobic Ile111 and Phe104 residues creating the binding site for **c)** the helicase-interacting motif HIM1 and **d)** the HIM2 of the gp61 primase. Interaction surfaces of HIM1 and HIM2 are colored based on the level of the hydrophobicity from dark cyan to wheat (inset).

**Supplementary Figure 10.**
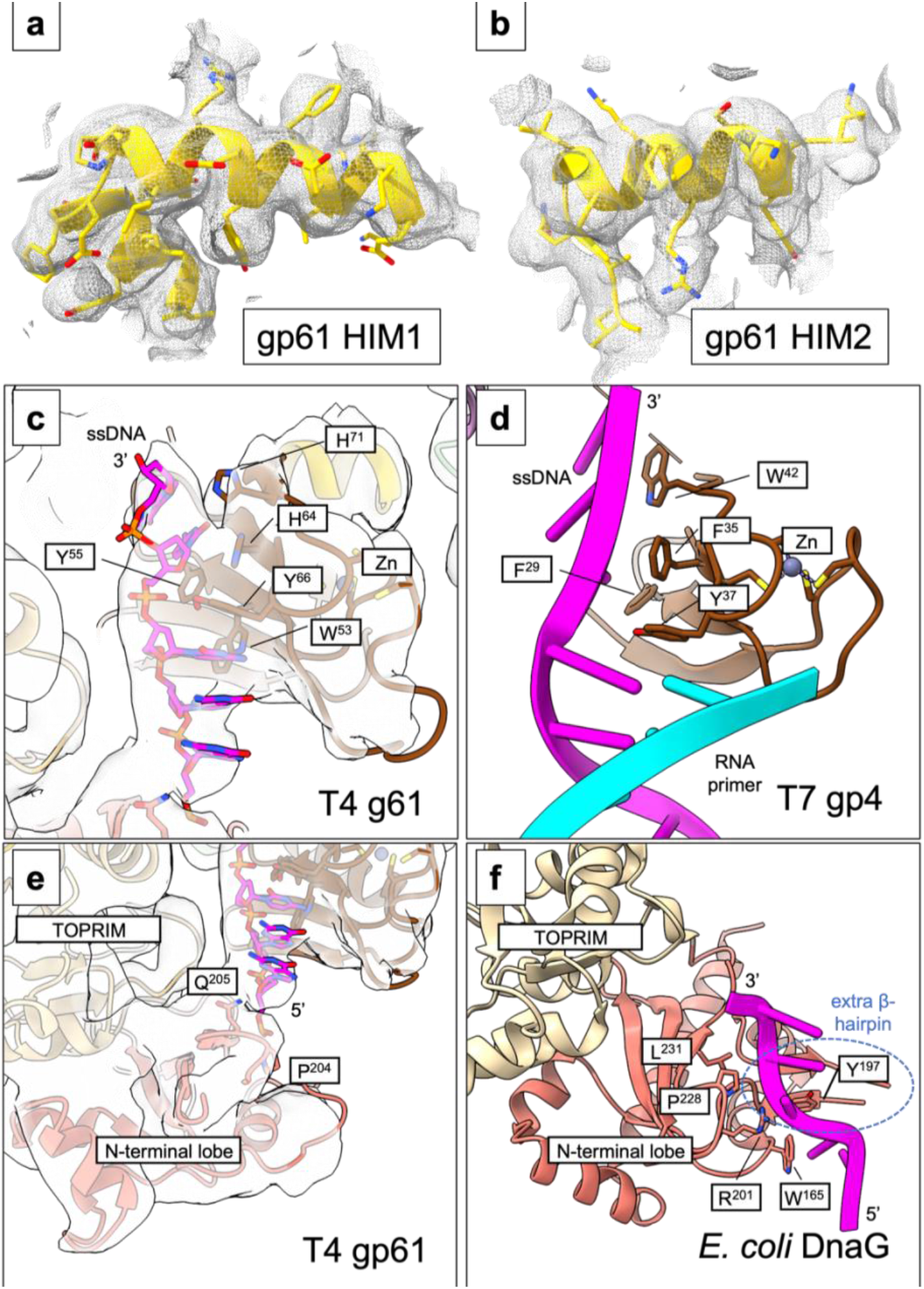
Local EM densities of the gp61 primase. EM densities of **a)** gp61 HIM1 and **b)** gp61 HIM2. Comparison of the interactions between the conserved ZBD and the ssDNA in **c)** the gp61 primase (this study, primosome pose 1) and **d)** in the T7 gp4 (PDB 6N9U). The ssDNA interacts with the conserved residues Tyr66 in the gp61 primase and Tyr37 in the gp4 primase. Comparison of the interaction between the RPD and the ssDNA in **e)** the gp61 primase (this study, primosome pose 1) and **f)** the *E. coli* DnaG primase (PDB 3B39). In both structures, the ssDNA interacts with the β-hairpin in the RPD N-terminal lobe. We note that the DnaG primase has an additional β-hairpin.

**Supplementary Figure 11.**
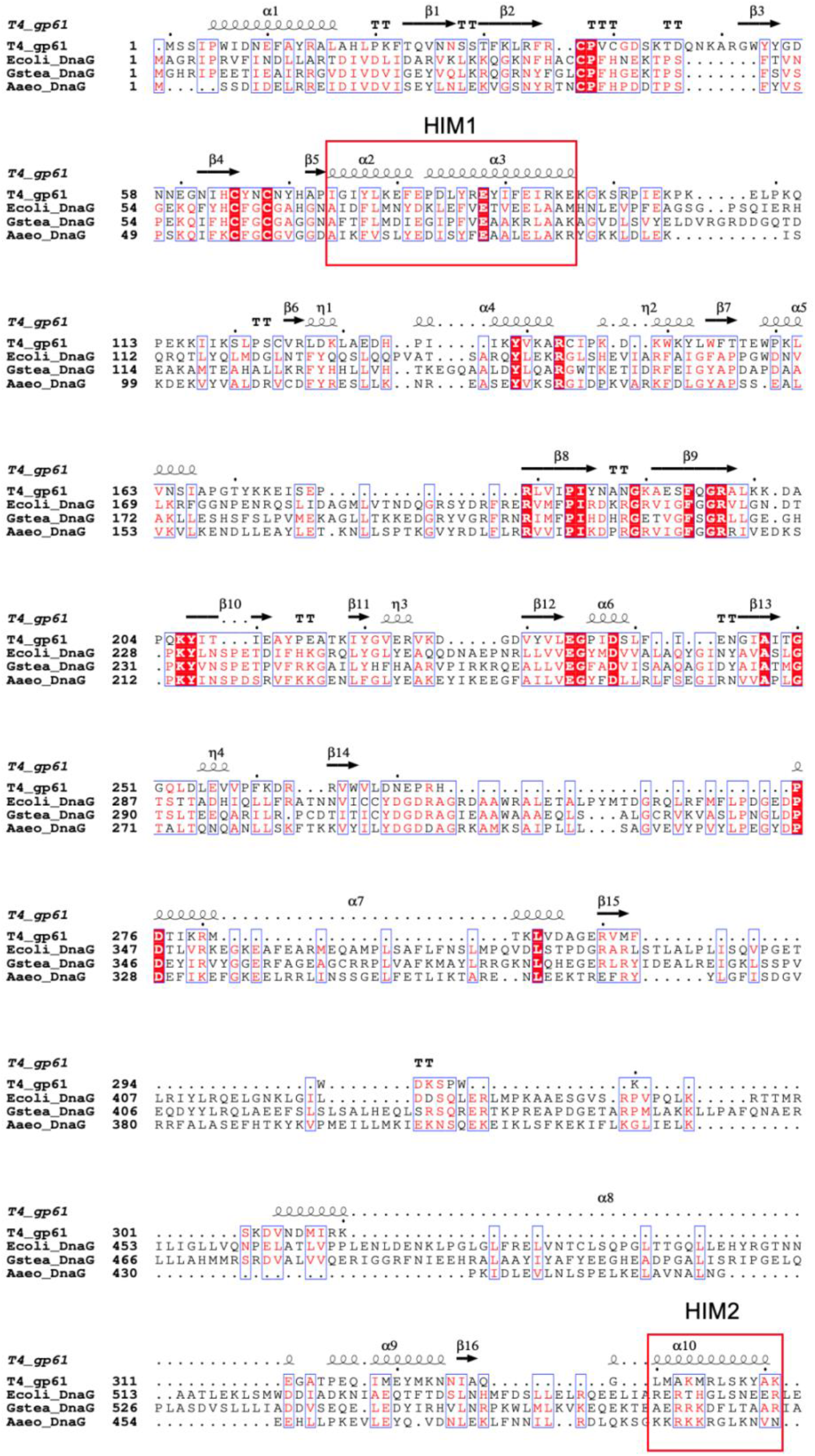
Sequence alignment of the T4 gp61 with several bacterial DnaG primases. The helix-interacting motifs HIM1 and HIM2 of gp61 are highlighted by red boxes in the sequence alignment demonstrating that the interfacial residues of Pro83, Arg87, Ile90, Phe91, and Arg94 are not conserved in the DnaG primases. Ecoli: *Escherichia coli*; Gstea: *Geobacillus stearothermophilus*; Aaeo: *Aquifex aeolicus*. Note that the gp61 HIM2 structurally aligns with the C-terminal HID of the DnaG primases, but their sequences are not conserved.

**Supplementary Figure 12.**
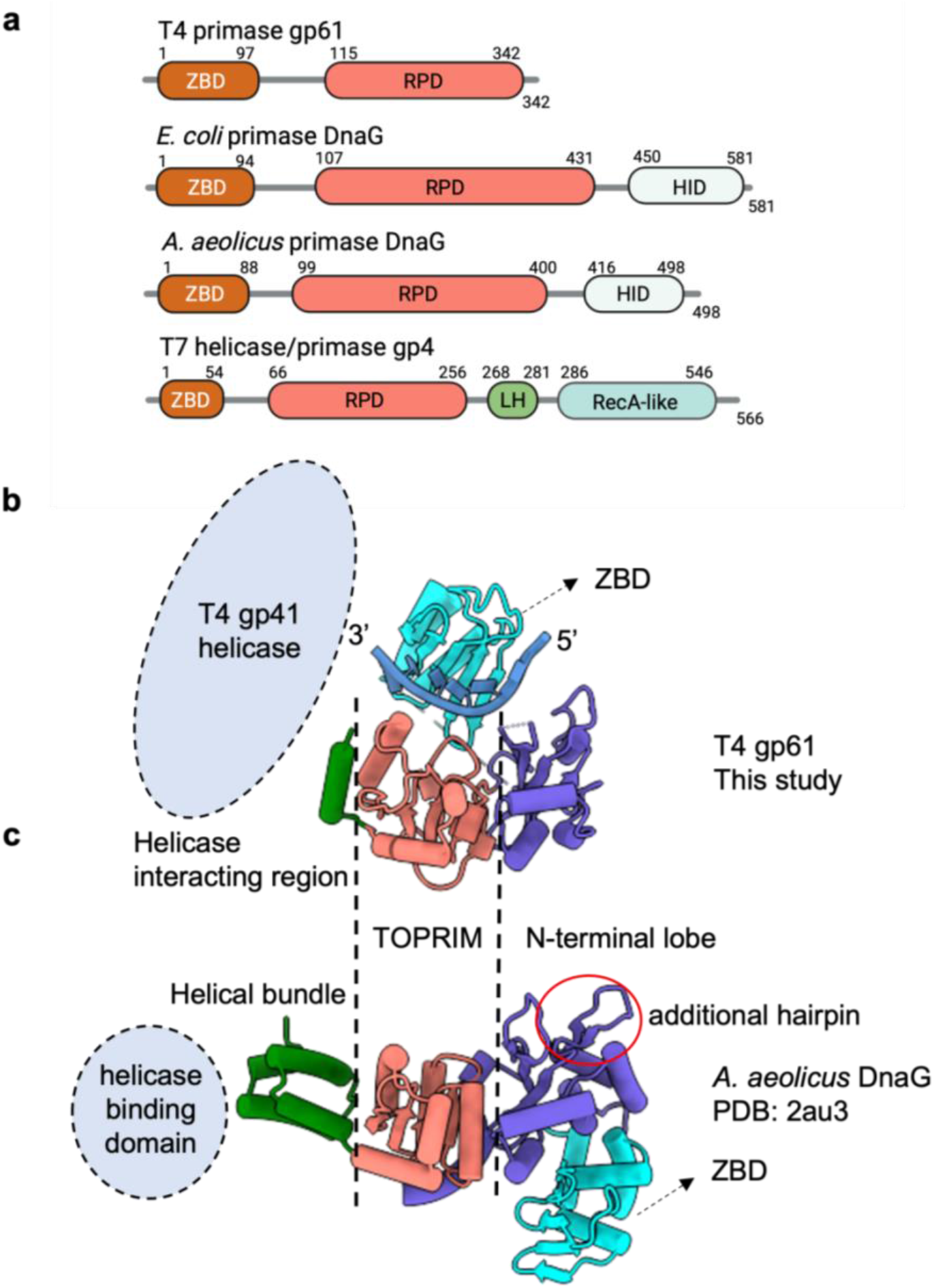
Structural comparison of the T4 gp61 and the bacterial DnaG primases. **a)**Domain architecture of the T4 gp61, T7 gp4, and two bacterial DnaG primases. The structures of **b)** the gp61 primase (this study) and **c)** the DNA-free *A. aeolicus* DnaG primase (PDB 2AU3) are aligned and shown separately for clarity. The TORPIM domain is similar in both structures, and the DnaG helicase interaction domain (HID) functions similarly to the gp61 HIM2. The N-terminal lobe of the DnaG RPD contains an ssDNA-binding β-hairpin (red circle) that is absent in the gp61 primase. The location of the ZBD is different between the primases; the ssDNA-binding surfaces of the gp61 RPD and ZBD face the ssDNA.

**Supplementary Figure 13.**
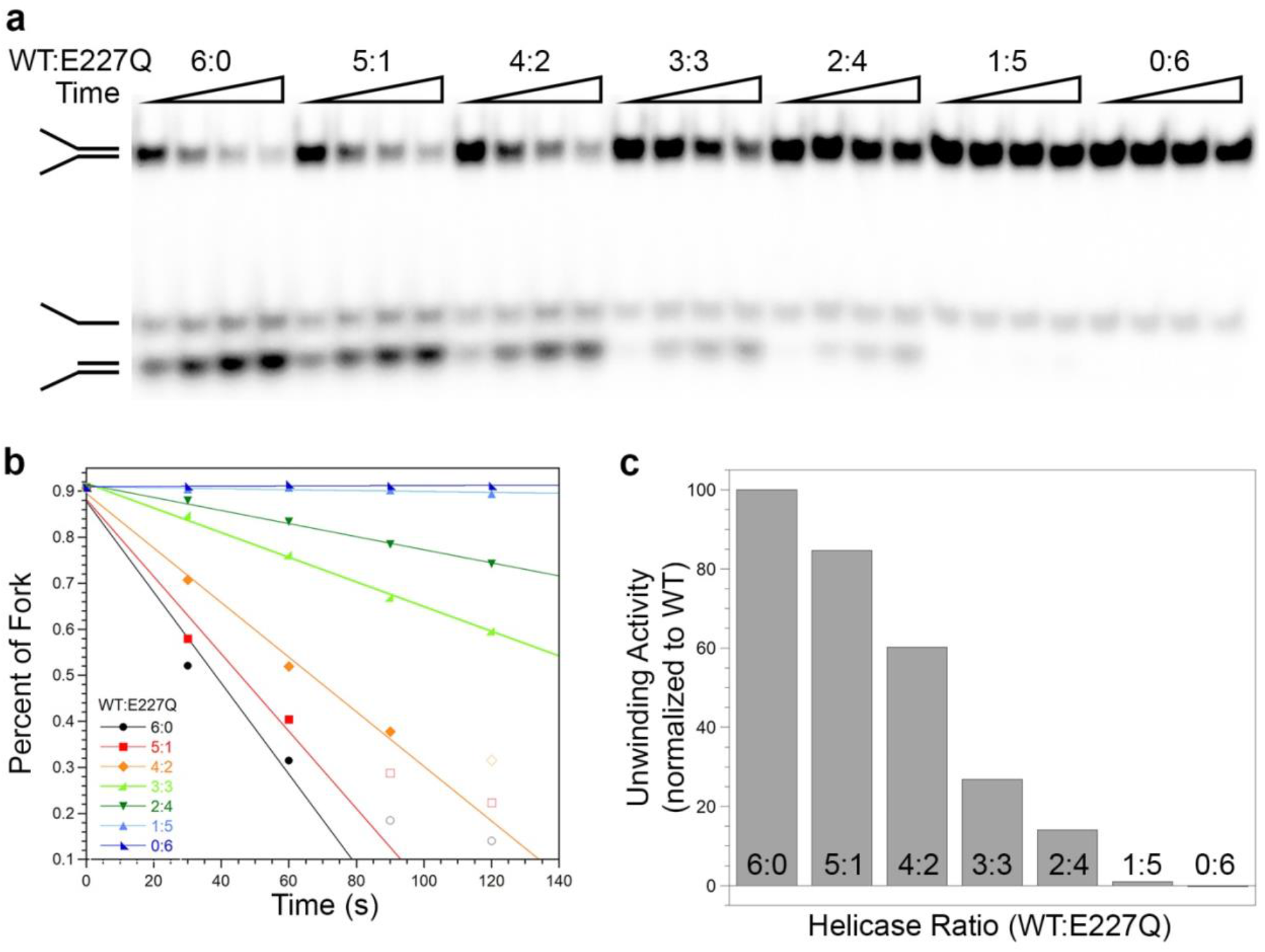
Unwinding activity of the gp41(E227Q) helicase mutant. **a)**Native polyacrylamide gel analysis of the unwinding activity of various ratios of WT gp41: gp41(E227Q) totaling 300 nM helicase at time 30, 60, 90, and 120 s. **b)** The unwinding rate for each ratio is calculated from the plot of the quantified percentage of fork DNA substrate versus time. **c)** Bar graph illustrating the loss of unwinding activity as the ratio of the gp41(E227Q) helicase mutant is increased until there is no unwinding activity with only gp41(E227Q) helicase present in the reaction.

**Supplementary Figure 14.**
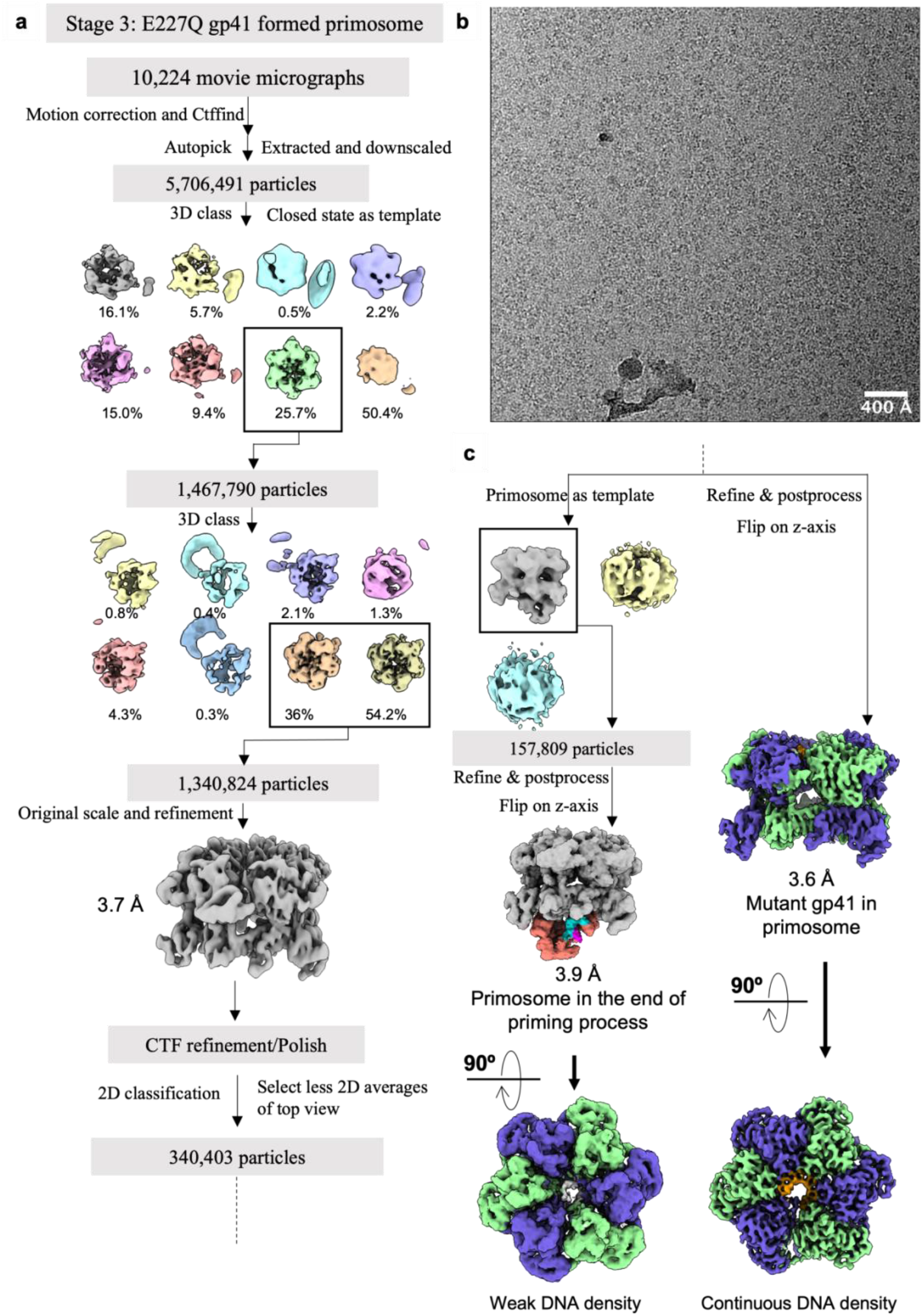
Processing of the cryo-EM images from the mutant T4 primosome assembly sample. **a)**Data processing flowchart. **b)** A typical raw micrograph. **c)** Additional data processing steps leading to the EM maps of the gp41 helicase(E227Q) alone (right) and the mutant primosome consisting of the gp41 helicase(E227Q) and the gp61 primase (left).

**Supplementary Figure 15.**
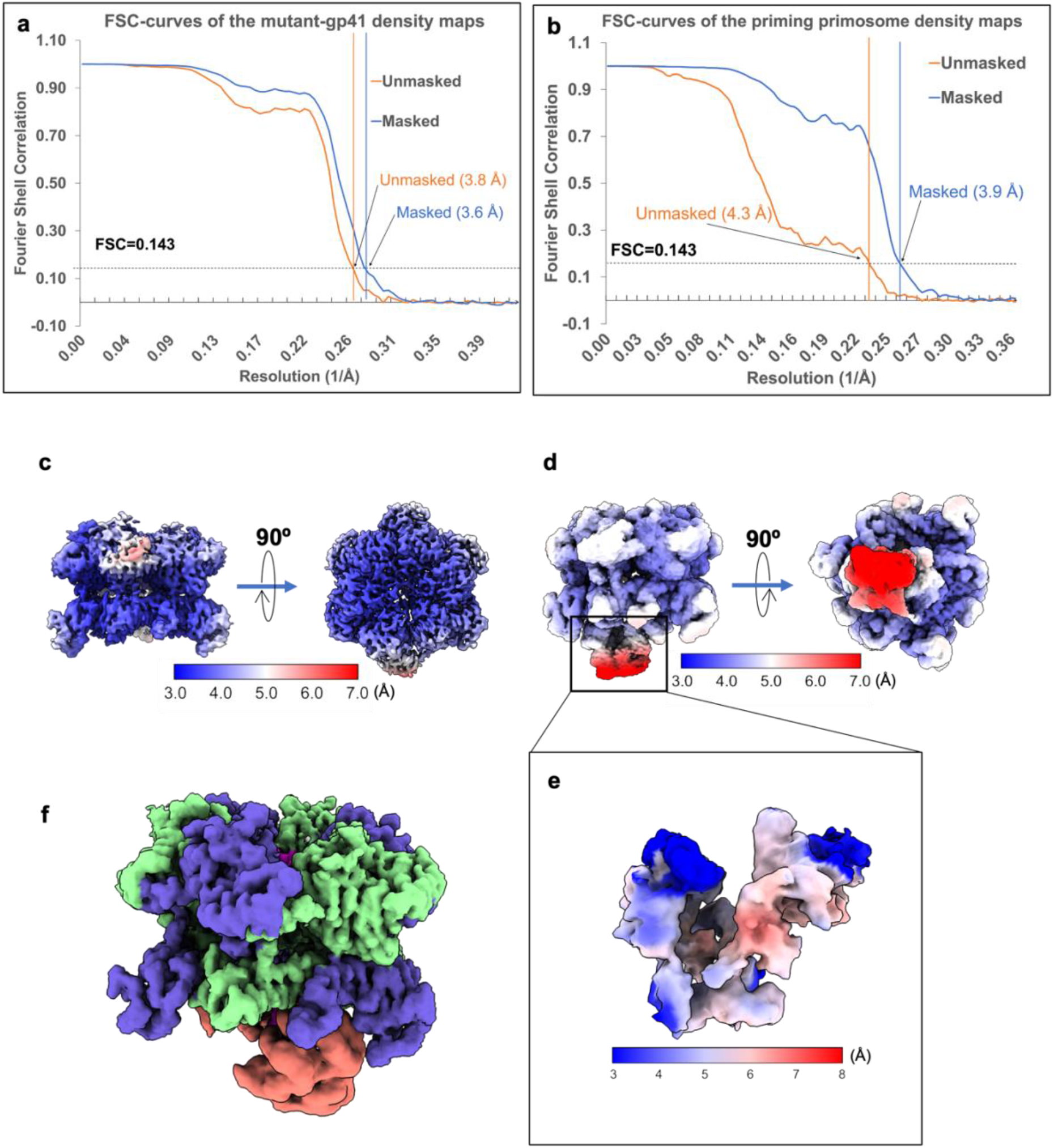
Resolution estimation of the mutant T4 primosome EM maps. FSC curves of the 3D EM maps of **a)** the ssDNA-bound gp41 helicase(E227Q) and **b)** the mutant primosome. The mutant primosome consists of a well-defined primase region and averaged helicase region, indicating multiple binding orientations of the primase. Local resolution maps of **c)** the ssDNA-bound gp41 helicase(E227Q), **d)** the mutant primosome, and **e)** the focus refined primase region. **f)** The focus refined primase and helicase were combined into a composite map and colored by chain.

**Supplementary Figure 16.**
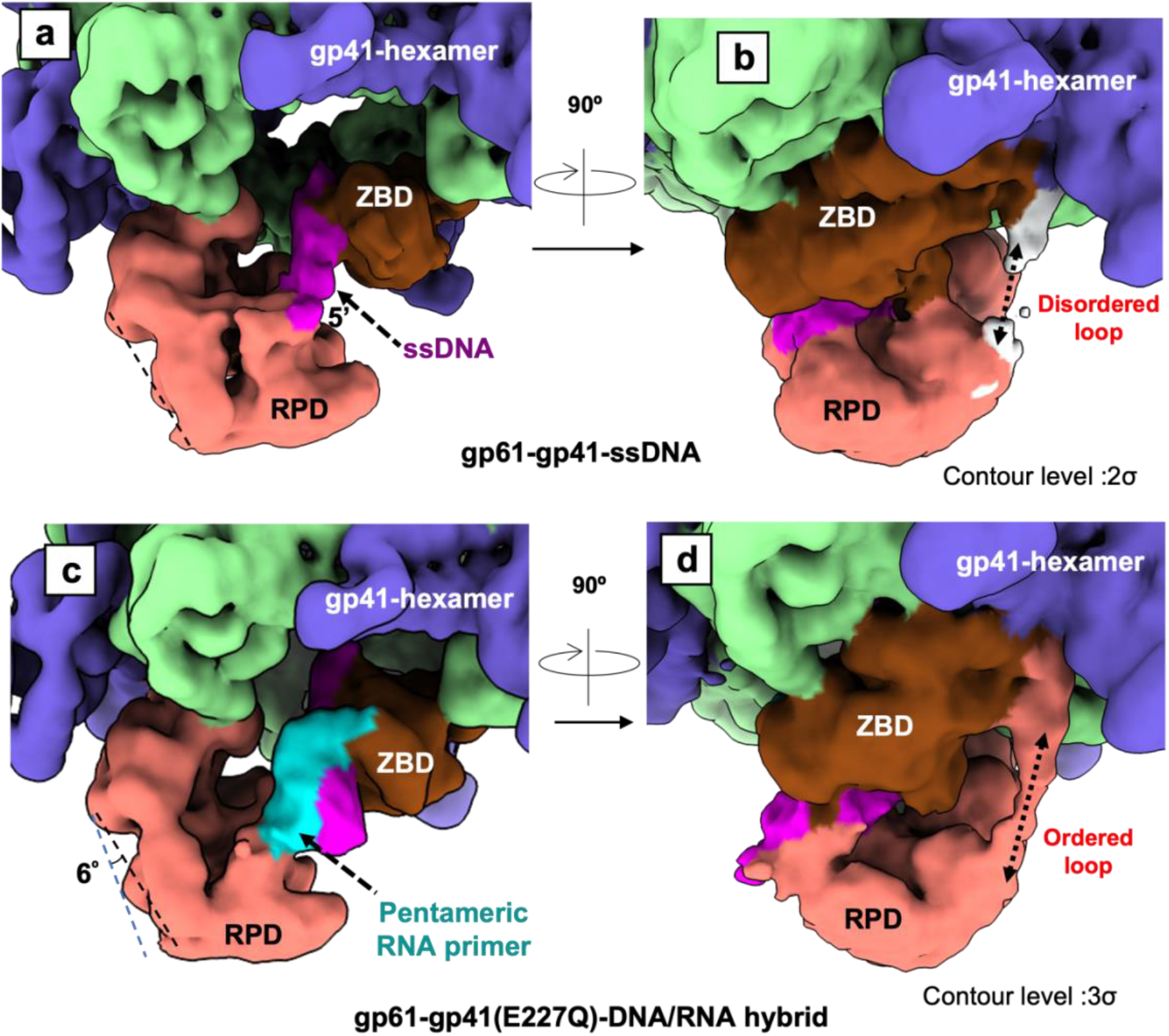
Comparison of the EM maps of the T4 primosome in potential DNA-scanning and post RNA primer-synthesis modes. **a-b**) Two views of the EM map of the WT T4 primosome likely in a DNA-scanning mode. **c-d**) Two views of the EM map of the mutant T4 primosome likely in a post RNA primer-synthesis mode. Note that the gp61 linker loop connecting the ZBD and RPD is disordered in **b**) the potential DNA-scanning mode, but becomes well ordered in **d**), likely a post RNA primer-synthesis mode.

**Supplementary Figure 17.**
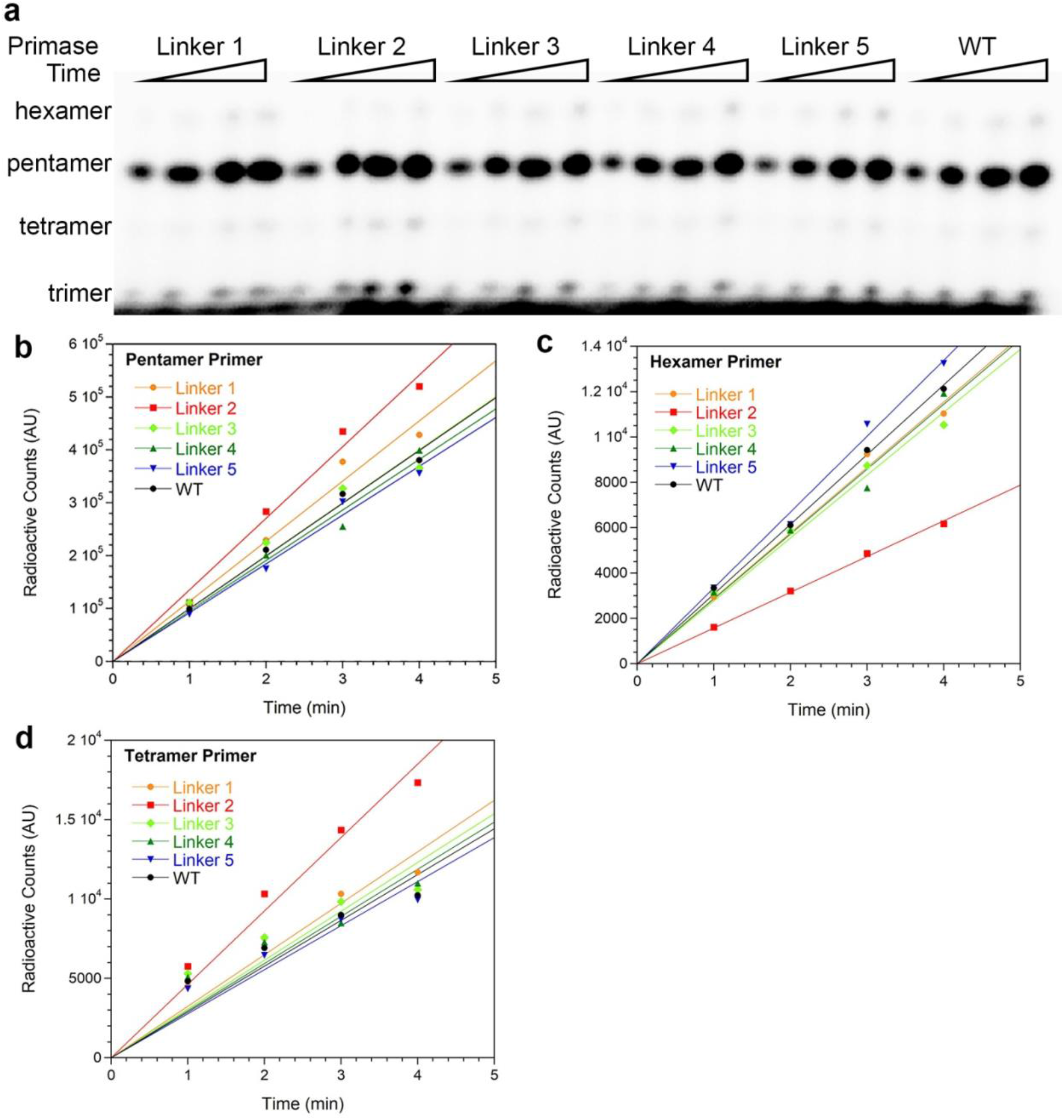
Priming activity of the series of gp61 primase linker loop mutants. **a)**Denaturing polyacrylamide gel analysis of the priming activity of the series of gp61 primase linker loop mutants at time 1, 2, 3, and 4 min. The priming rate for each gp61 primase linker loop mutant is calculated from the plot of the quantified amount of **b)** pentameric, **c)** hexameric, or **d)** tetrameric RNA primer synthesized versus time. While trimeric RNA primer products are observed on the polyacrylamide gel, their quantification is unreliable due to the lack of complete separation from free radiolabeled CTP.

**Supplementary Video 1. The assembly process of the T4 primosome.** The video starts with the inactive open-spiral gp41 helicase, morphs to the ssDNA template bound and still inactive helicase, then transitions to the active closed-ring helicase with a large conformational change, and finally shows the gp61 primase binding to the active helicase hexamer; this completes the assembly of the T4 primosome.

**Supplementary Video 2. The two functional states of the T4 primosome.** The video begins by showing the morph among the three binding poses of the gp61 primase on the gp41 helicase hexamer in the primosome state 1, which is suggested to be a DNA-scanning mode. Then, the scene transitions to focus on the primase binding region, morphing between states 1 and 2. State 2 is likely a post RNA primer-synthesis state. Transition from state 1 to 2 is repeated twice to highlight the rotation of the RPD primase domain. The dashed line represents the linker loop between the ZBD and RPD and is suggested to constrain the RPD rotation range and influence the length of the synthesized primer.

## Notes

### Competing Interest Statement

The authors have declared no competing interest.

